# Fused in sarcoma regulates DNA replication timing and progression

**DOI:** 10.1101/2020.04.22.055343

**Authors:** Weiyan Jia, Mark A. Scalf, Peter Tonzi, Robert J. Millikin, Sang Hwa Kim, William M. Guns, Lu Liu, Adam S. Mastrocola, Lloyd M. Smith, Tony T. Huang, Randal S. Tibbetts

## Abstract

*Fused in sarcoma (FUS*) encodes a low complexity RNA-binding protein with diverse roles in transcriptional activation and RNA processing. While oncogenic fusions of FUS and transcription factor DNA-binding domains are associated with soft tissue sarcomas, dominant mutations in FUS cause amyotrophic lateral sclerosis (ALS) and frontotemporal dementia (FTD). FUS has also been implicated in DNA double-strand break repair (DSBR) and genome maintenance. However, the underlying mechanisms are unknown. Here we employed quantitative proteomics, transcriptomics, and DNA copy number analysis (Sort-Seq), in conjunction with *FUS^-/-^* cells to ascertain roles of FUS in genome protection. *FUS^-/-^* cells exhibited alterations in the recruitment and retention of DSBR factors BRCA1 and 53BP1 but were not overtly sensitive to genotoxins. By contrast, FUS-deficient cells had reduced proliferative potential that correlated with reduced replication fork speed, diminished loading of pre-replication complexes, and attenuated expression of S-phase associated genes. FUS interacted with lagging strand DNA synthesis factors and other replisome components, but did not translocate with active replication forks. Using a Sort-Seq workflow, we show that FUS contributes to genome-wide control of DNA replication timing and is essential for the early replication of transcriptionally active DNA. These findings illuminate new roles for FUS in DNA replication initiation and timing that may contribute to genome instability and functional defects in cells harboring disease-associated FUS fusions.

## Introduction

Fused in sarcoma (FUS, also referred to as translocated in liposarcoma, TLS) is a member of the FET (FUS, EWSR1, TAF15) family of RNA- and DNA-binding proteins that play important roles in transcription and splicing (Shang and Huang, 2016; Tan and Manley, 2009). Originally described as an oncogenic fusion to the CHOP transcription factor in myxoid liposarcoma (MLS) (Crozat et al., 1993; Rabbitts et al., 1993), *FUS* rose to prominence with the discovery that inherited and de novo mutations in its open-reading frame cause dominant forms of amyotrophic lateral sclerosis (ALS) and frontotemporal dementia (FTD) (Corrado et al., 2010; Kwiatkowski et al., 2009; Vance et al., 2009). Although the underlying mechanisms are still unclear, the preponderance of ALS/FTD-associated mutations in FUS interfere with its nuclear import and folding, leading to the accumulation of cytosolic FUS aggregates that disrupt cellular function through loss- and gain-of function mechanisms impacting protein translation and nuclear transport among other processes (Dormann et al., 2010; Kamelgarn et al., 2018; Ling et al., 2019; Lopez-Erauskin et al., 2018; Shang and Huang, 2016).

FET proteins share a common domain structure that includes an N-terminal low-complexity domain (LCD), a Gly-rich region, one or more arginine/glycine-rich (RGG) domains, an RNA recognition motif (RRM) with RNA- and DNA-binding activity, a zinc-finger domain, and a carboxyl-terminal PY-type nuclear localization signal that interacts with transportin nuclear import receptors that are essential for proper FUS folding (Dormann et al., 2010; Hofweber et al., 2018; Qamar et al., 2018; Tan and Manley, 2009; Yoshizawa et al., 2018). The LCD is also of particular interest as it exhibits strong transcriptional coactivation potential *in vitro* and the fusion of this domain to the CHOP DNA binding domain drives gene deregulation and oncogenesis in MLS (Tan and Manley, 2009; Zinszner et al., 1994). The LCD also mediates protein-protein interactions and participates in FUS oligomerization and liquid demixing (Han et al., 2012; Kwon et al., 2013; Qamar et al., 2018; Schwartz et al., 2013; Sun et al., 2011) that may be central to its normal roles in transcription and splicing and pathologic roles in ALS/FTD (Shang and Huang, 2016).

In addition to their accepted roles in RNA processing, several lines of evidence support a role for the FET proteins in the cellular DNA damage response (DDR). FUS participation in the DDR was first inferred from chromosome instability and mild radiosensitive phenotypes of *FUS^-/-^* mice (Bertrand et al., 1999; Hicks et al., 2000; Kuroda et al., 2000). FET proteins are capable of promoting invasion and pairing of a homologous ssDNA sequence with a dsDNA molecule *in vitro* (Baechtold 1999; Bertrand 1999; Guipaud 2006), which suggests a possible role for FET proteins in the D-loop formation step of homology-directed DNA double-strand break repair (HDR), while other studies showed that the FUS LCD is phosphorylated in response to DNA damage by DNA damage-activated protein kinases DNA-PKcs and ATM (Gardiner et al., 2008; Han et al., 2012), which are important regulators of non-homologous end joining. Consistent with a direct or indirect role for FUS in DSBR, we and others showed that shRNA-mediated depletion of FUS reduced the repair of HDR and NHEJ reporter substrates (Mastrocola et al., 2013; Rulten et al., 2014; Wang et al., 2013).

A role in the DNA damage response is further suggested by poly (ADP)-ribosyl (PAR) polymerase (PARP)-dependent localization of FUS to sites of microirradiation-induced DNA damage (Mastrocola et al., 2013; Rulten et al., 2014; Wang et al., 2013). FUS is capable of interacting directly with PAR chains through its RGG-domain (Mastrocola et al., 2013) and the FET proteins are heavily PARylated in response to genotoxic stress (Jungmichel et al., 2013). Mechanistically it was reported that FUS mediates the recruitment of histone deacetylase 1 (HDAC1), KU70, NBS1, and phosphorylated H2AX (γH2AX) and ATM at sites of DNA damage and that this recruitment pathway as well as FUS-dependent repair was compromised by ALS/FTD-associated mutations (Wang et al., 2013). It has also been proposed that FUS organizes DSBs in a PARP-dependent manner for their subsequent repair (Singatulina et al., 2019); while Wang et al. reported that FUS recruits DNA ligase III downstream of PARP activation to repair SSBs and that ALS-associated mutations in FUS disrupt its SSB repair activity (Wang et al., 2018). Finally, it was recently reported that FUS regulates the response to transcription-associated rDNA damage via association with Topoisomerase 1 in the nucleolus (Martinez-Macias et al., 2019). Despite these studies, the molecular mechanisms linking FUS to the different repair pathways in which it has been implicated remain unclear and the extent to which FUS-dependent RNA processing may contribute to reported DDR phenotypes in FUS-deficient cells is not known.

Here, we probed FUS-dependent genome protection using transcriptomic, proteomic, and functional analysis of *FUS^-/-^* cell lines. Our findings suggest that genome protection functions of FUS are particularly important in S-phase, where it contributes to BRCA1 recruitment, replicon initiation, and replication timing. These studies provide new insights into FUS-mediated genome protection in mitotically active cells.

## Results

### Generation and phenotypic characterization of *FUS^-/-^*cells

To discern roles of FUS in genome protection we disrupted *FUS* gene loci in U-2 OS osteosarcoma cells using CRISPR/CAS9 followed by functional reconstitution with an untagged FUS retroviral expression (see Materials and methods). To ensure rigorous results we studied multiple *FUS^-/-^* clones and selected a reconstituted *FUS^-/-^:FUS* line with physiologic levels of FUS expression (Fig. 1A-C). Notably, protein levels of TAF15 and EWSR1 were not upregulated in *FUS^-/-^* U-2 OS cells, diminishing concerns about functional compensation.

**Figure 1.**
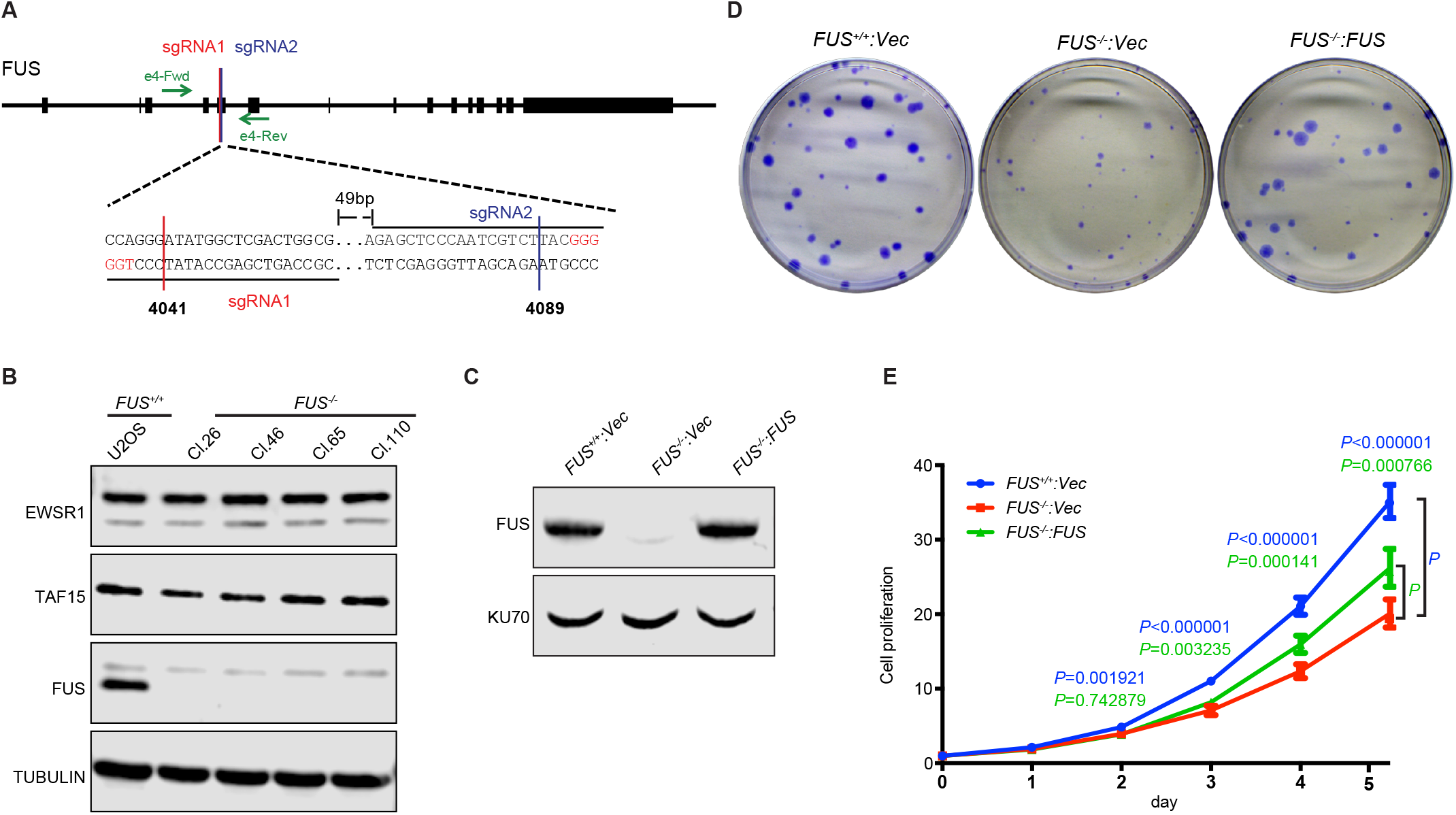
FUS promotes cell proliferation. A, Schematic of the FUS gene targeting. Two guide RNAs, sgRNA1 and sgRNA2, were used to target FUS exon 4 (See methods). The primers, e4-Fwd and e4-Rev, were used for genotyping. B, Expression of FET proteins (FUS, EWSR1, and TAF15) in *FUS^-/-^* clones. Note lack of compensation by EWSR1 and TAF15. C, Reconstitution of *FUS^-/-^* (Cl.110) with an untagged FUS retroviral vector. The same vector expressing β-Glucuronidase was introduced as a negative control into *FUS^-/-^* cells. D, *FUS^-/-^* cell colonies exhibited reduced growth relative to *FUS^+/+^* and *FUS^-/-^*:*FUS* cells. E, Cell proliferation rates of *FUS^+/+^*, *FUS^-/-^* and *FUS^-/-^*: *FUS* U-2 OS cells. Each sample contains 6 technical replicates. The bars represent mean ± SD. The*T* test was performed and the *P* values were shown on plot.

FUS knockdown cells displayed mild IR sensitivity and modest defects in the repair of NHEJ and HDR reporter substrates (Mastrocola et al., 2013) while a second study reported that that FUS knockdown suppressed γH2AX and 53BP1 focus formation (Wang et al., 2013). We assessed time courses of γH2AX and 53BP1 accumulation and dissolution at IR-induced foci in *FUS^+/+^*, *FUS^-/-^*, and *FUS^-/-^*:*FUS* U-2 OS cells exposed to 2 Gy IR. The initial recruitment of γH2AX and 53BP1 to IRIF was comparable between *FUS^+/+^* and *FUS^-/-^* cell lines; however, 53BP1 and γH2AX foci persisted longer in *FUS^-/-^* cells relative to *FUS^+/+^* cells (Sup. Fig. 1A-C). Similar findings were made for RIF1,which functions downstream of 53BP1 to promote NHEJ (not shown). Prolonged accumulation of γH2AX and 53BP1 at IR-induced foci may reflect persistent DSBs.

In contrast to findings for 53BP1, the frequency of cells displaying IR-induced BRCA1 foci was reduced in *FUS^-/-^* cells, and this was corrected by FUS reexpression (Sup. Fig. 2 A-D). The defect in BRCA1 focus formation was specific to conditions of DNA damage since unirradiated *FUS^-/-^* cells displayed more numerous BRCA1 foci than unirradiated *FUS^+/+^* cells (Sup. Fig. 2C). Given its upstream role in DNA end resection and HDR(Chen et al., 2018a), reduced recruitment of BRCA1 may contribute to modest HDR defects seen in *FUS^-/-^* cells (Mastrocola et al., 2013). Despite the changes in 53BP1 and BRCA1 recruitment to IR-induced foci, *FUS^-/-^* cells did not exhibit significant hypersensitivity to mechanistically distinct genotoxins, including hydroxyurea (HU, replication stress), mitomycin C (MMC, DNA crosslinker), camptothecin (CPT, Top1 inhibitor), and calicheamicin γ1 (CLM, radiomimetic) (Sup. Fig. 3). These findings suggest FUS is not a core mediator of DSBR in U-2 OS cells.

### *FUS^-/-^* cells exhibit defects in DNA replication

*FUS^-/-^* U-2 OS cells exhibited reduced colony outgrowth and proliferative potential that was corrected by FUS reexpression (Fig. 1D and E). Proliferation and colony growth defects were seen across independent *FUS^-/-^* close as well as *FUS^-/-^* H460 lung adenocarcinoma cells (not shown). Following synchronous release from G_1_/S phase arrest, *FUS^-/-^* cells exhibited reduced reentry and progression through S phase, which was particularly pronounced at the 6 h timepoint (Fig. 2A). The S-phase delay in *FUS^-/-^* cells was further revealed through EdU incorporation experiments. Specifically, *FUS^-/-^* cells exhibited reduce S-phase entry 6 h following release from a double thymidine block and accumulated in G_2_/M to a lesser degree than *FUS^+/+^* or *FUS^-/-^:FUS* cells 12 h following release (Fig. 2B and Sup. Fig. 4). These experiments also revealed slightly reduced levels of EdU incorporation in asynchronously growing *FUS^-/-^* cells relative to *FUS^+/+^* or *FUS^-/-^:FUS* cells (Fig. 2B).

**Figure 2.**
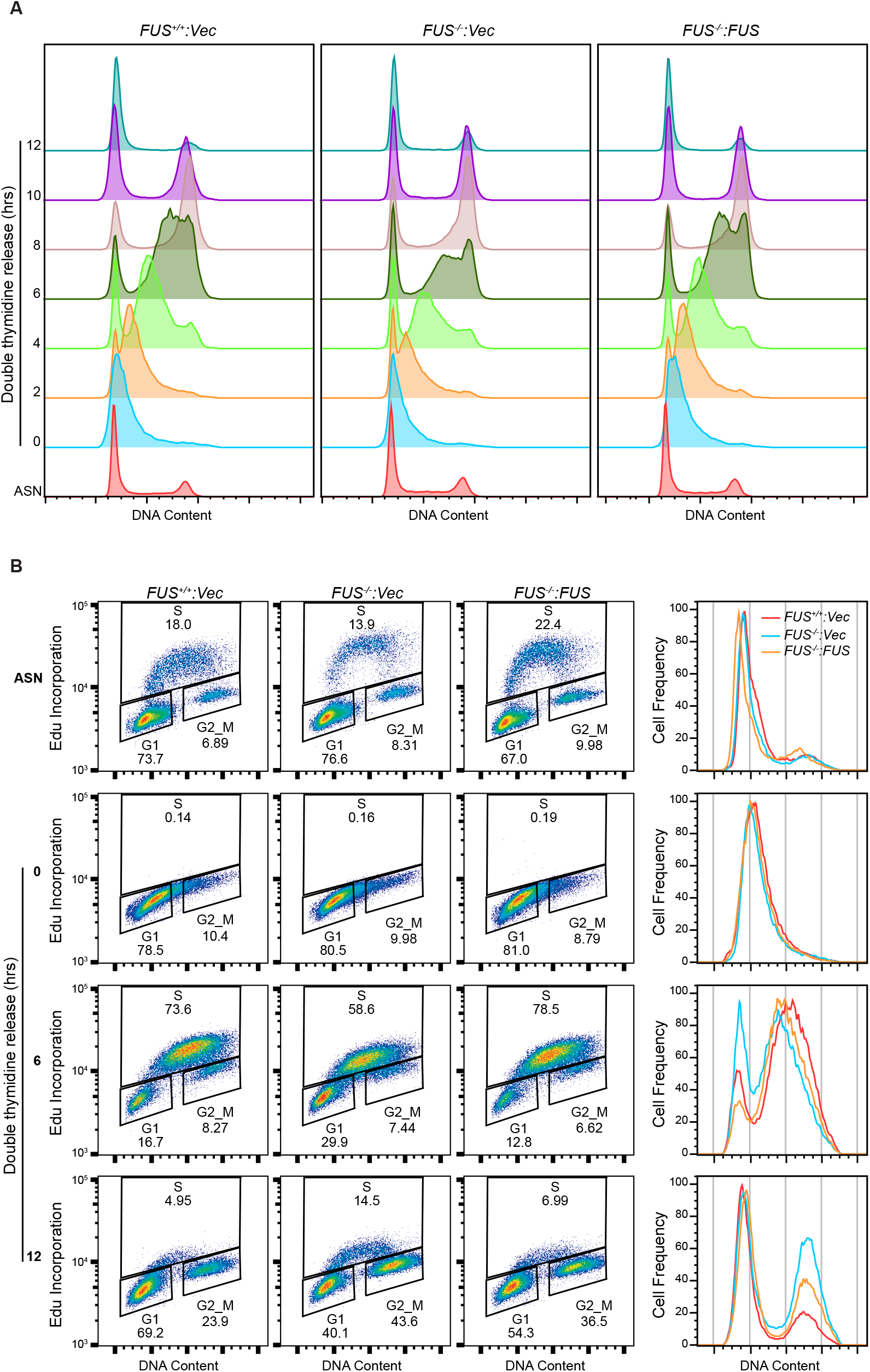
FUS is required for S-phase progression. A, DNA replication progression was analyzed by PI staining and flow cytometry. Cells were synchronized to G_1_/S boundary by double thymidine block and released into fresh growth medium for the indicated times and stained with PI for cell cycle analysis. B, DNA progression was monitored by EdU incorporation under the same conditions as in (A). additional timepoints are presented in Sup. Fig. 4.

We performed RNA-Seq to establish gene expression correlates for DNA replication defects of FUS-deficient cells. We identified 626 genes that were differentially expressed between *FUS^+/+^* and *FUS^-/-^* cells that were corrected by FUS reexpression (Sup. Fig. 5A and B, Sup. Tab. 3 and 4). Gene set enrichment analysis(GSEA) revealed that cell cycle, DNA repair and DNA replication processes were downregulated while immunomodulatory pathways were upregulated in in *FUS^-/-^* cells (Sup. Fig. 5C, Fig. 3A and 3D). DNA replication-associated genes that were downregulated in *FUS^-/-^* cells included *GINS4, MCM4*, and *RFC3, RCF4*, and *TIMELESS*, (Fig. 3B and C). DNA repair related genes, including *WRN, PRKDC, FANCD2, FANCA, RAD52*, were also downregulated in *FUS^-/-^* cells (Fig. 3E and F). Interestingly, the NHEJ factor *53BP1* was upregulated in *FUS^-/-^* cells (Sup. Fig. 5E). A subset of gene expression changes evident in RNA-Seq data were confirmed by qPCR (Sup. Fig. 5D and E). In sum, downregulation of S-phase genes is compatible with reduced proliferative potential of *FUS^-/-^* cells.

**Figure 3.**
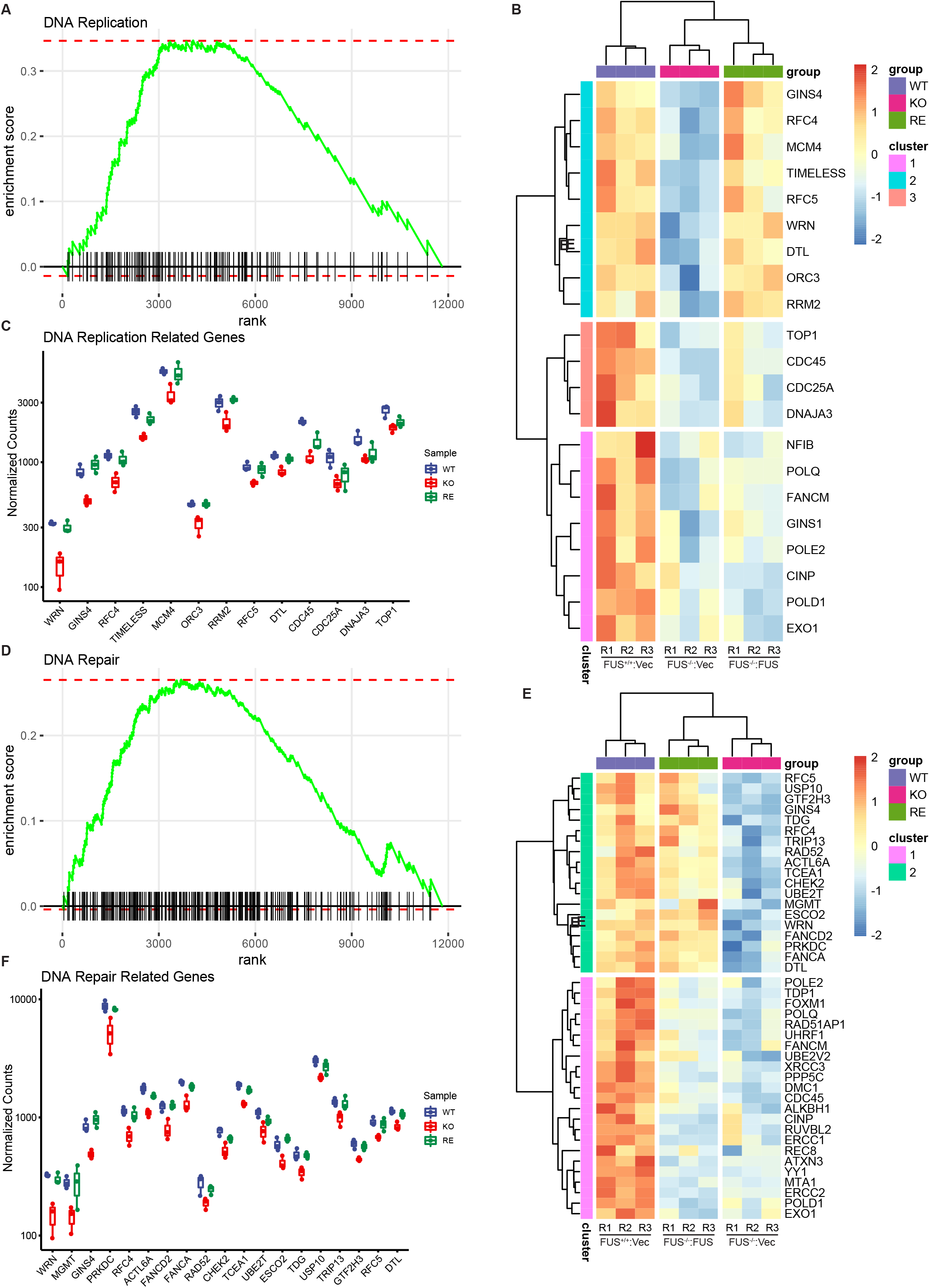
Reduced expression of replication-associated genes in FUS deficient cells. A, Enrichment plot of DNA replication pathway from GSEA analysis using GO gene sets (biological process) in the Sup. Tab. 3. B, Heatmap of differentially expressed DNA replication genes. Genes were clustered to three groups based on ward.D2 method. C, Normalized RNA-Seq counts of cluster 2 genes involved in the DNA replication pathway. D, Enrichment plot of DNA repair pathway from GSEA analysis using GO gene sets (biological process) in the in the Sup. Tab. 3. E. Heatmap of the leading gene list of DNA repair pathway shown significate changing in all samples. Genes were clustered into two groups based on ward.D2 method. F, DNA repair related gene expressions of cluster 1 were shown in normalized counts from RNA sequencing results.

To ascertain impacts of FUS deficiency on replication fork (RF) dynamics, we performed DNA fiber analysis (Tonzi et al., 2018), on *FUS^-/-^*, *FUS^+/+^*, *FUS^-/-^*:*FUS* cells sequentially labeled with IdU and CldU. *FUS^-/-^* cells exhibited significant reductions in CldU track lengths indicative of reduced DNA replication rate (Fig. 4B). *FUS^-/-^* cells also showed delayed RF restart following release from a transient hydroxyurea (HU) block (Fig. 4C). Both replication velocity and replication restart phenotypes were rescued by FUS reexpression. Because a reduced rate of DNA replication can lead to micronucleus formation and genomic instability(Hoffelder et al., 2004), we measured micronuclei in *FUS^+/+^*, *FUS^-/-^*, and *FUS^-/-^*:*FUS* U-2 OS cells treated with a low dose of the DNA polymerase alpha inhibitor aphidicolin. *FUS^-/-^* cells exhibited increased rates of micronucleus formation relative to *FUS^+/+^* and *FUS^-/-^*:*FUS* U-2 OS cells (Fig. 4A), suggesting that FUS enhances genome stability under replication stress.

**Figure 4.**
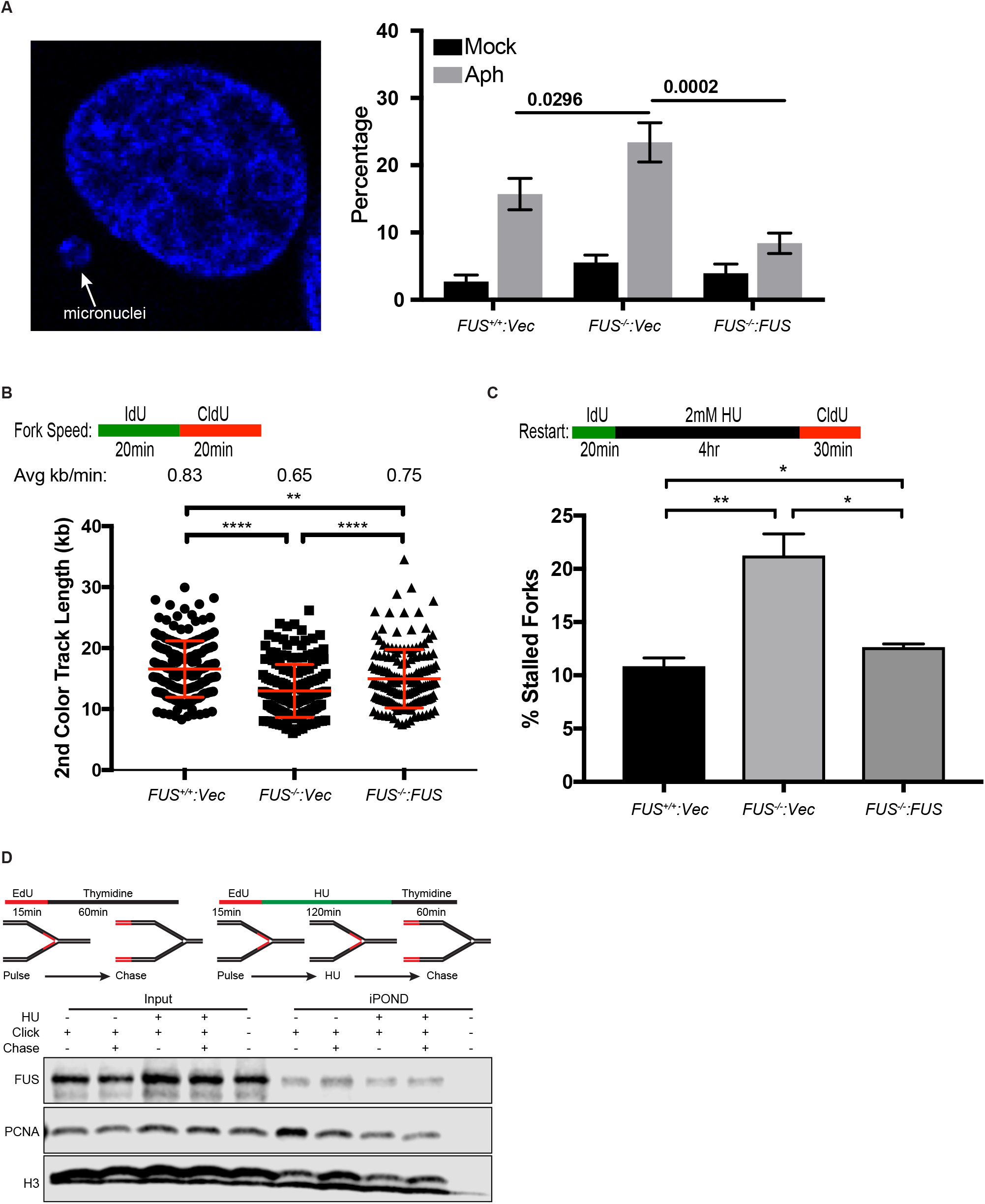
FUS deficiency leads to genomic instability and replication stress. A, *FUS^+/+^*, *FUS^-/-^* and *FUS^-/-^*: *FUS* U-2 OS cells were treated with or without 0.2 μM aphidicolin (Aph) for 24 hours, fixed and stained with DAPI for micronucleus counting. *P* values were calculated by two-way ANNOVA test. Data are means ± SE(n=3 biological replicates). More than 250 cells for each sample in each biological replicate were counted. B, Replication fork speed is reduced in *FUS^-/-^* cells. The second pulse (CIdU) was used for measurement of track length, which was converted to micrometers using a 1 μm =2.59 kb conversion factor. The average fork length was divided by 20 min to derive replication speed. C, Replication fork restart was measured as shown in the schematic. Percentages of fork restart (% stalled forks) in HU-treated cells are shown. Data are mean± s.d.(n=3). *P* values were calculated using a t-test with Welch’s correction. n.s. = no significance, * = p <0.05, ** = p <0.01, *** = p <0.001, **** = p <0.0001. D, FUS does not translocate with the replisome. An iPOND assay was performed as shown in the schematic. HEK 293T cells were pulse labeled with 20 μM EdU for 15 min and then chased with 20 μM thymidine for 1 h. For replication stress, cells were treated with 2 mM HU after EdU labeling, and then chased with thymidine for another 1 hour. Western blotting was used to assess FUS, PCNA, and histone H3 enrichment at EdU-labeled replication forks.

### FUS regulates pre-replication complex (pre-RC) loading and associates with DNA replication factors

Given their reduced DNA replication rate, we investigated whether *FUS^-/-^* cells exhibited defects in the chromatin loading of replication licensing factors, including the origin recognition complex (ORC), CDC6, CDT, and the MCM replicative helicase (Fragkos et al., 2015). Mitotically arrested *FUS^+/+^*, *FUS^-/-^*, and *FUS^-/-^*:*FUS* cells were released into early G_1_ phase and soluble and chromatin fractions analyzed by immunoblotting. FUS-deficient cells showed normal cell progression from G_2_/M to G_1_ phase and unchanged ORC loading onto chromatin in G_1_ (Fig. 5A and B). By contrast, recruitment of CDC6 and CDT1 was significantly decreased in *FUS^-/-^* cells and rescued by FUS reexpression (Fig. 5B). As expected, CDC6 and CDT1-dependent loading of MCM complex was also reduced in *FUS^-/-^* cells (Sup. Fig. 6). Collectively, these results revealed that FUS facilitates ORC-dependent recruitment of pre-RC factors CDC6 and CDT1 to replication origins.

**Figure 5.**
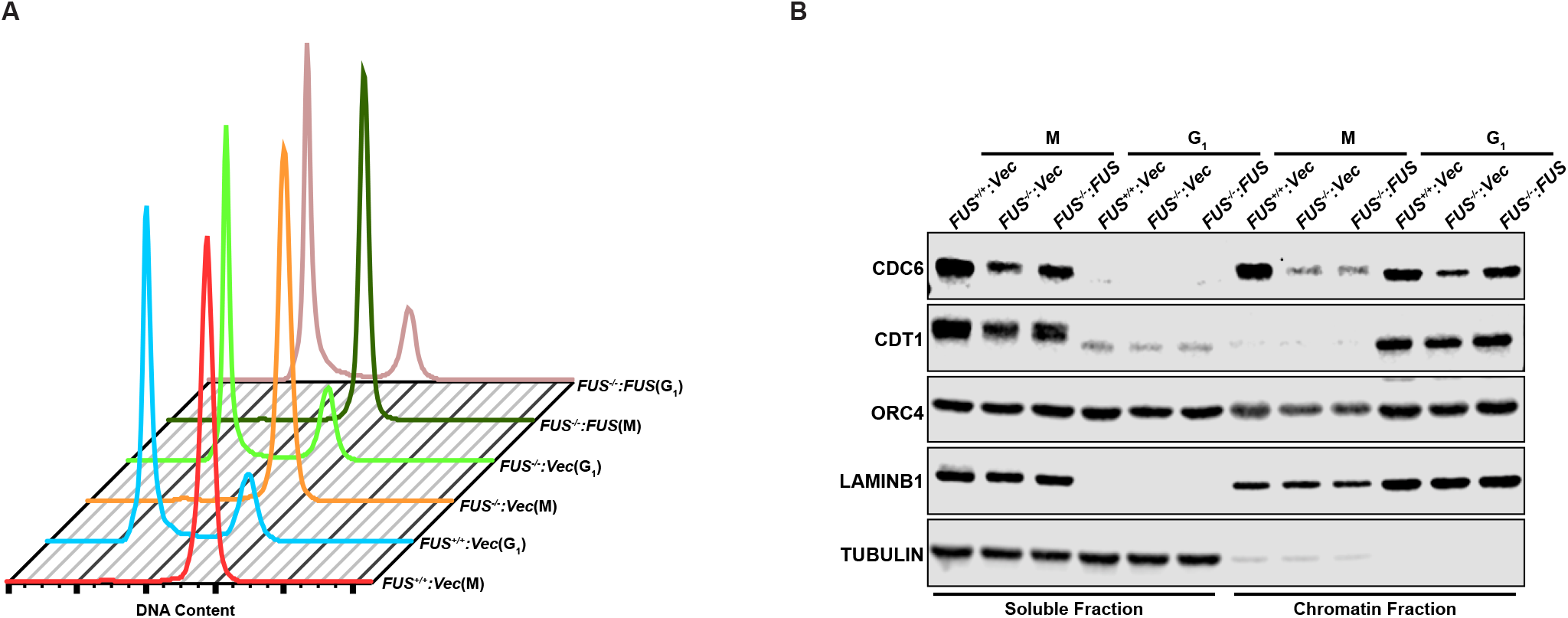
FUS regulates pre-replication complex(pre-RC) loading onto chromatin. A, Cell cycle profiles of *FUS^+/+^*, *FUS^-/-^*, and *FUS^-/-^*: *FUS* U-2 OS cells that were synchronized in early M phase with nocodazole and then harvested or released into G_1_ phase for 5 h. B, Chromatin loading of ORC and pre-RC proteins in *FUS^+/+^*, *FUS^-/-^* and *FUS^-/-^*: *FUS* U-2 OS cells. Mitotic and G_1_ fractions were immunoblotted with the indicated antibodies.

Reasoning that FUS may play direct roles pre-RC loading and or DNA repair, we performed quantitative proteomic analysis of FUS complexes using a chromatin-IP procedure in which endogenous FUS-chromatin complexes were digested with nuclease prior to immunoprecipitation with α-FUS antibodies and analysis by quantitative mass spectrometry (MS) (Mohammed et al., 2016). The same chromatin-IP procedure was carried out using *FUS^-/-^* cells as a negative control. GSEA using all identified FUS interactants revealed RNA processing, DNA repair, and DNA replication as functional processes that were statistically overrepresented in the dataset of FUS-interacting proteins (Sup. Fig. 7A and Sup. Tab. 1 and 2). The abundance of RNA-binding proteins (RBPs) in FUS complexes is consistent with other published studies (Kawaguchi et al., 2020; Sun et al., 2015). Nucleotide excision repair and DNA strand elongation proteins were among the most significantly enriched pathways within the DNA replication/repair gene sets (Sup. Fig. 7A). We plotted those proteins within DNA repair and replication GO terms that showed a nominal 1.3-fold enrichment in IPs from *FUS^+/+^* cells relative to *FUS^-/-^* cells (Fig. 6A). Proteins of interest include DSBR factors (DNA-PK, Ku70, Ku80, PNKP), SSBR/BER proteins (PARP1, FEN1, PNKP, APEX1), DNA replication factors (DNA polymerase δ (POLδ or POLD1), PCNA, and UHRF1), and topoisomerases (TOP1, TOP2α). The presence of SSBR/BER factors, including PARP is consistent with the ability of FUS to bind to PAR chains (Mastrocola et al., 2013) while the presence of POLδ but not POLε in FUS IPs is interesting given their participation in leading strand and lagging strand DNA synthesis, respectively (Burgers and Kunkel, 2017). We carried out validation co-IP assays to confirm that endogenous FUS interacted with TOP1, PCNA, POLδ1, and FEN1 in unsynchronized (Fig 6B) or synchronized S phase cells (Fig 6C) and further validated association between FUS and POLδ1, PCNA, and FEN1 in proximity ligation (PLA) assays (Fig. 6D and E).

**Figure 6.**
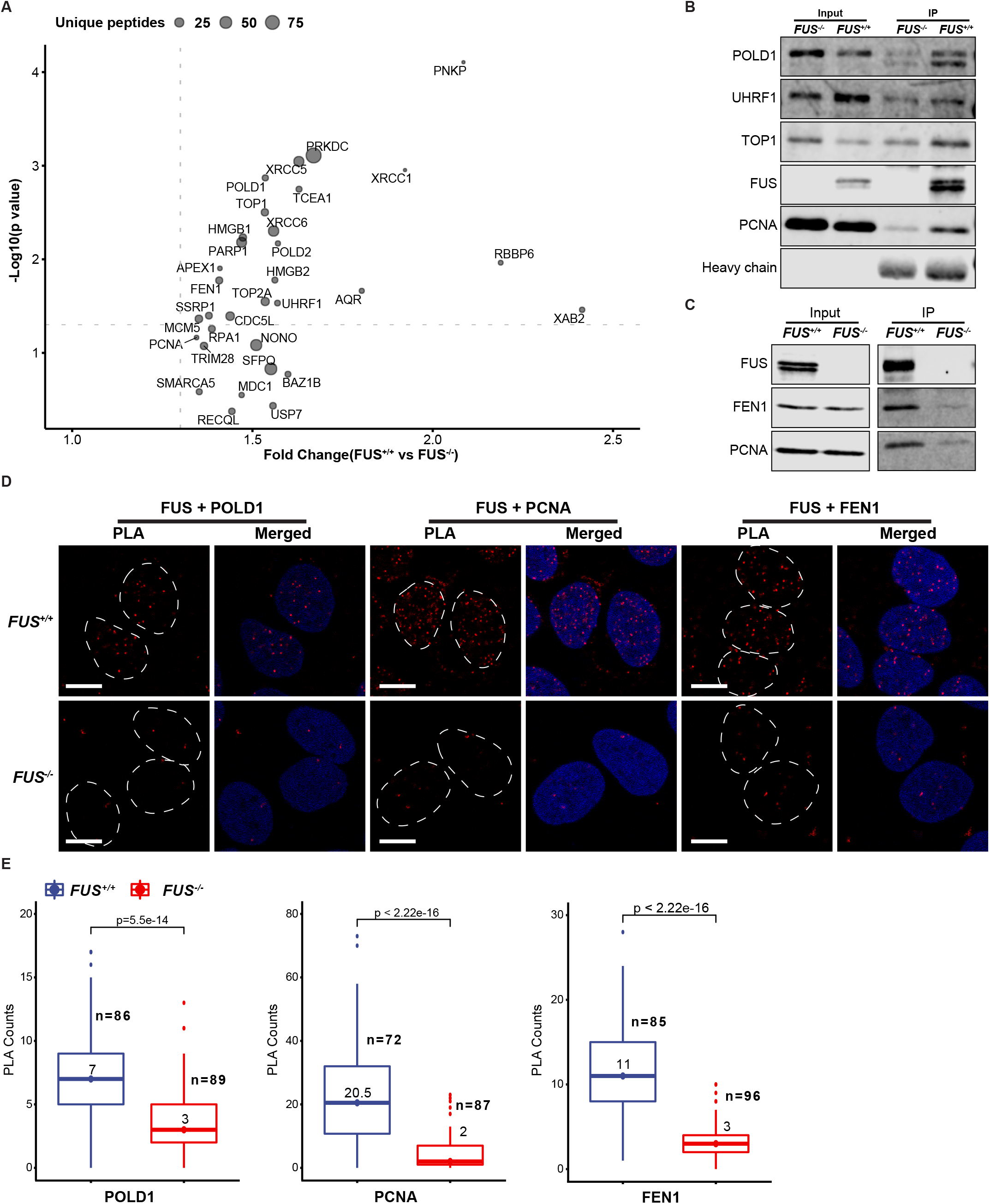
FUS interacts with DNA repair and DNA replication factors. A, FUS interacting proteins were identified by cross-linking chromatin immunoprecipitation (IP) and analyzed by MS. The results are combination of three biological replicates quantified by non-isotopic spectral peptide counting. The data shown are DNA repair (red colored) and DNA replication (blue colored) pathway related interactions based on GSEA (full listed shown in Sup. Fig. 7A). The unique peptides are summarized from three replicates raw data. The grey doted lines are 1.3 of fold change and 0.05 of *P* value. B, Coimmunoprecipitation of FUS with POLD1, UHRF1, TOP1 and PCNA in unsynchronized cells. C, Coimmunoprecipitation of FUS with FEN1 and PCNA in synchronized S phase cells. D, In situ proximity ligation (PLA) assay was employed to verify the interactions between FUS, and POLD1, PCNA and FEN1. Nuclear regions were cycled by dashed lines in PLA red channel based on DAPI signal. E, Quantification results of PLA signal in (D). The values are median of PLA foci in each sample. *P* values were calculated by Wilcox test method.

The replication defects in *FUS^-/-^* cells and interaction with DNA replication factors raised the possibility that FUS directly participates in DNA replication. To investigate this possibility, we carried out an iPOND assay that measures the association of proteins with nascently synthesized DNA (Sirbu et al., 2011). *FUS^+/+^*, *FUS^-/-^* and *FUS^-/-^*:*FUS* U-2 OS were pulse labeled with EdU and then chased with thymidine in the absence or presence of 2 mM HU prior to formaldehyde crosslinking and isolation of EdU-protein complexes. As expected, the abundance of the PCNA sliding clamp in EdU-labeled complexes decreased during the thymidine chase period as the replisome advanced beyond the region of nascent, EdU-labeled DNA (Fig. 3D). Although FUS was also observed in EdU-labeled DNA, its abundance was slightly increased following thymidine chase, as was histone H3 (Fig. 3D). A similar iPOND labeling pattern has been described for DNA-binding proteins such as HMGA1 and LaminB1 that maintain high-order chromatin (Lopez-Contreras et al., 2013). This result suggests that FUS is proximal to replication factors on chromatin, but does not translocate with the active replisome.

### FUS regulates DNA replication timing

Chromosomal replication is stochastically initiated from hundreds of origins that fire with characteristic, heritable timing (Fragkos et al., 2015). Replication timing (RT) can be qualitatively evaluated according to the pattern of 5-Bromo-2’-deoxyuridine (BrdU) or 5-Ethynyl-2’-deoxyuridine (EdU) incorporation following synchronized release from a double thymidine block (Dimitrova and Berezney, 2002). Early S-phase cells exhibit a uniform EdU incorporation pattern (Fig. 7A, white arrows); middle S-phase cells exhibit perinuclear and perinucleoloar EdU incorporation (Fig. 7A, yellow arrows); and late S-phase cells exhibit large puncta of EdU incorporation (Fig. 7A, green arrows). Origins with shared firing kinetics are topologically organized into chromatin subdomains in a process that requires RIF1 (Cornacchia et al., 2012; Dileep et al., 2015; Kanoh et al., 2015; Mattarocci et al., 2016; Sima et al., 2018; Yamazaki et al., 2012); however, few other timing regulators have been identified.

**Figure 7.**
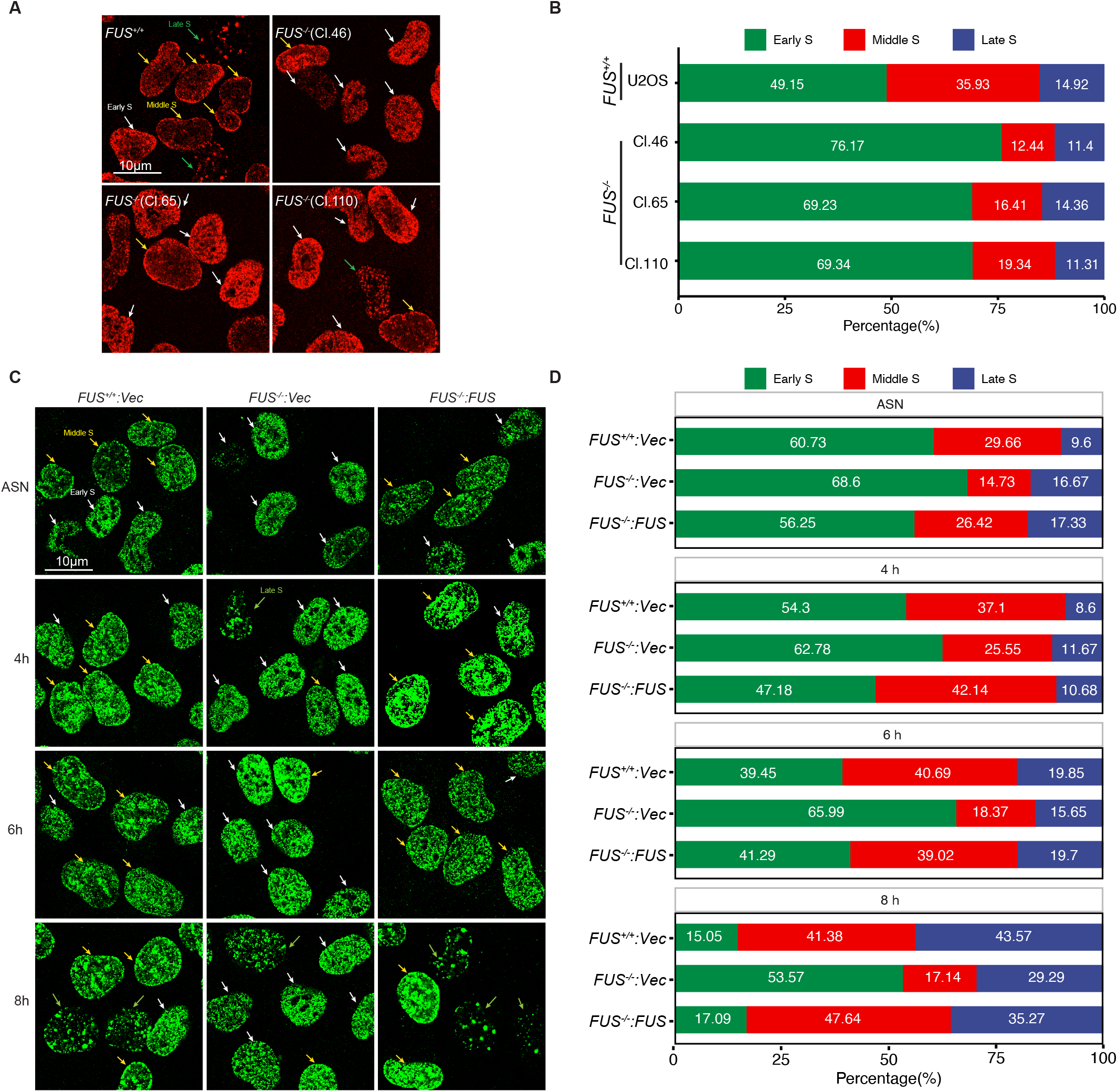
FUS regulates DNA replication timing (RT). A, Asynchronous U-2 OS cells and three *FUS^-/-^* clones (Cl. 46, Cl. 65 and Cl. 110) were pulse-labeled with EdU for 20 min and scored for the presence of early, mid, or late EdU staining patterns. B, Quantification analysis of cell numbers of each S phase patterns in (A), and the percentage were calculated in each sample. C, Cells were synchronized with double thymidine and then released into S phase for indicated times. Cells were then pulse-labeled with BrdU, stained and imaged by confocal microscopy. D, Quantification results of samples using a minimum 100 cells per sample (C).

We noted that the frequency of mid-S phase staining patterns was reduced ~50% in *FUS^-/-^* cells relative to *FUS^+/+^* cells (Fig. 7A and B), suggesting a potential RT defect. To rule out the apparent defect was not due to delayed S-phase entry of *FUS^-/-^* cells we carried out a timecourse analysis of *FUS^+/+^*, *FUS^-/-^*, and *FUS^-/-^*:*FUS* cells released from thymidine block for 4, 6, or 8 h. *FUS^-/-^* cells exhibited reduced frequencies of the mid-S-phase staining pattern at all three timepoints, even though the frequency of late S-phase patterns more than doubled from 4-8 h (Fig. 7C and D). Importantly, FUS reexpression reversed the mid-S phase RT defect of *FUS^-/-^* cells (Fig. 7C and D). From this we conclude that FUS-deficient cells harbor RT defects that cannot be solely attributed to reduced rates of replication.

To follow up on the EdU labeling studies we measured genome-wide RT in *FUS^+/+^*, *FUS^-/-^* and *FUS^-/-^*:*FUS* U-2 OS cells using a Sort-Seq workflow (Koren et al., 2012). Propidium iodide-stained *FUS^-/-^*, *FUS^+/+^* and *FUS^-/-^*:*FUS* U-2 OS cells were sorted into G_1_ and S-phase fractions prior to genomic DNA isolation and deep sequencing (see Materials and Methods). The S/G_1_ phase read ratio was used to establish relative DNA copy between samples, with a higher ratio reflecting earlier RT (Fig. 8A). Using a fixed window method of read binning, we found that *FUS^-/-^* cells exhibited widespread changes in RT relative to *FUS^+/+^* and *FUS^-/-^*:*FUS* cells that was consistent across two biological replicates (Fig. 8C). FUS deficiency impacted bi-directional RT switches and was highly position dependent. For example, within the same 30 Mb interval of Chr18, *FUS^-/-^* cells exhibited accelerated RT (Fig. 8B, tan shading) and delayed RT (Fig. 8B, blue shading). The bi-directional RT switches in *FUS^-/-^* cells further suggests that timing changes in *FUS^-/-^* cells are not simply a byproduct of reduced RF speed.

**Figure 8.**
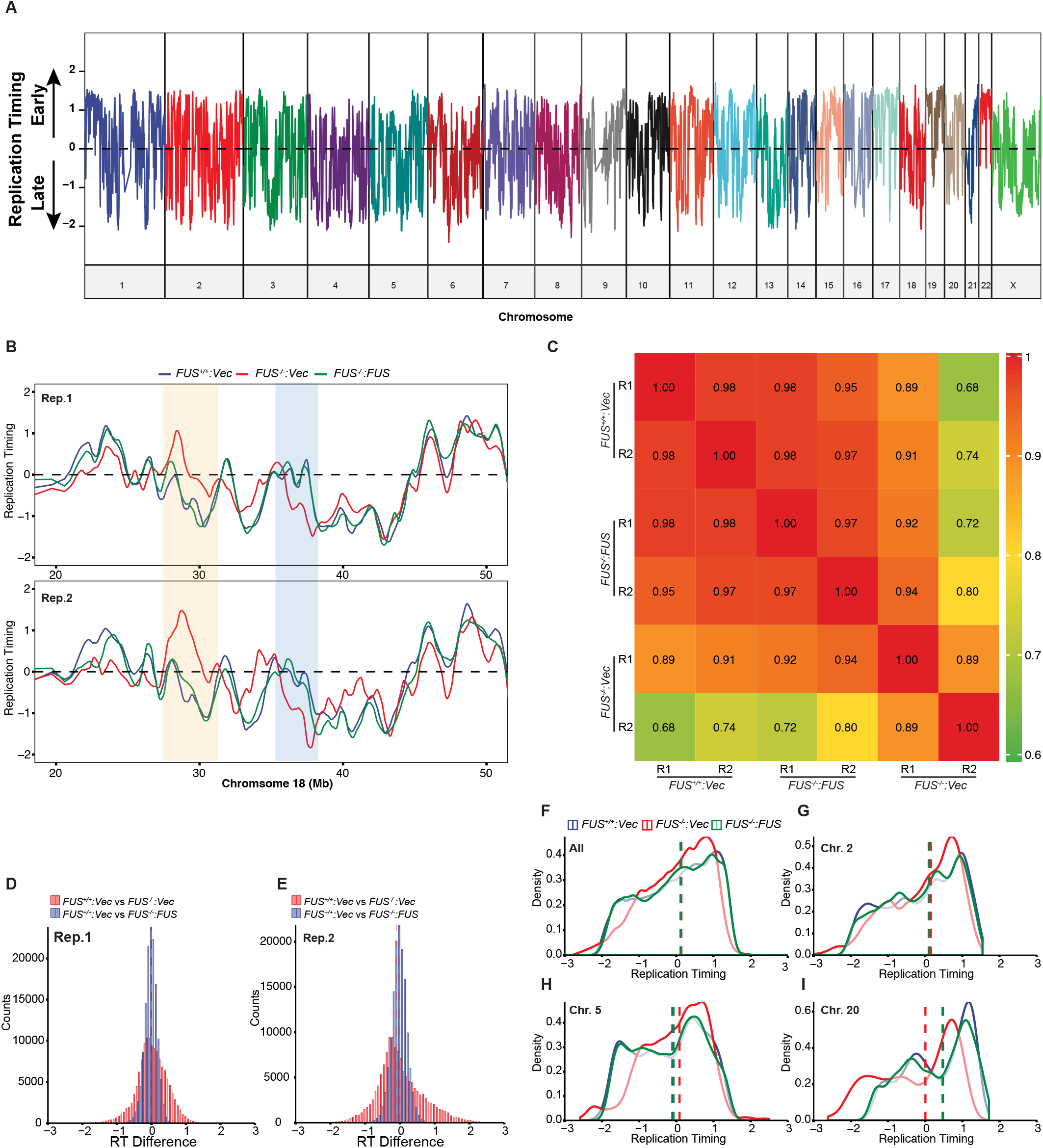
FUS influences genome wide RT. A, Whole genome wide replication timing profile of U-2 OS cells. The replication timing (RT) was calculated based on copy number variations between S and G_1_-phase cells (S/G_1_ ratio). The signal was normalized by Z-score and smoothed by Loess smoothing. B, Representative RT profiles of *FUS^+/+^*, *FUS^-/-^* and *FUS:FUS^-/-^* cells across two biological replicates. Regions of RT switching between *FUS^+/+^* and *FUS^-/-^* are highlighted. C, Correlation of RT between two biological replicates by Pearson’s method. The smoothed RT values were used for the correlation matrix. D, Genome-wide distribution of RT scores when comparing *FUS^+/+^* to *FUS^-/-^* or *FUS^+/+^* vs *FUS:FUS^-/-^* in two biological replicates. The bin sizes are 50 and 100 for Replicate1 (Rep.1) and Replicate2 (Rep.2) respectively. F to I, The RT density distribution for Rep. 2 was analyzed across all chromosomes (F), Chr.2 (G), Chr.5 (H) and Chr.20 (I). The RT density distribution for Rep. 1 are |shown in Sup. Fig. 8A to D. The dashed lines are the median of each sample. The Loess smoothed data was used for analysis.

Genome-wide bi-directional RT switches were further confirmed by RT distribution differences between *FUS^+/+^* and *FUS^-/-^* cells in both biological replicates (Fig.8D and E). Although the RT distribution of *FUS^-/-^* cells skewed slightly earlier than *FUS^+/+^* cells when examined across all chromosomes (Fig. 8F and Supp. Fig. 8A), RT directional changes were highly chromosome dependent. For example, while Chr2 did not show significant RT distribution differences between *FUS^+/+^* and *FUS^-/-^* cells, the RT distributions of Chr5 an Chr20 skewed early and late, respectively, in *FUS^-/-^* cells relative to *FUS^+/+^* cells (Fig. 8G to I and Sup. Fig. 8B to D). In summary, our data indicates FUS influences genome-wide RT in a chromosomal contextdependent manner.

### Characterization of FUS-associated replication domains

The above findings indicate that FUS deficiency leads to altered timing of chromosomal regions that we refer to as FUS-associated replication domains (FADs). To determine whether there are common features linking FADs we employed the unsupervised Segway deep learning tool (Chan et al., 2018; Hoffman et al., 2012) (Liu et al., 2016) to *de novo* segment replication domains (RDs) in our samples (see materials and methods). Three nonoverlapping contiguous segments were used to assign replication timing profiles into three types of RDs: early (ERD), late (LRD) and mid (MRD), which spans the transition between early and late zone. Genomic coverage of all three types of RDs did not significantly change between *FUS^-/-^* cells relative to *FUS^+/+^* cells (Fig. 9A and B). However, the average size of LRDs was significantly decreased in *FUS^-/-^* cells (Fig. 9C).

**Figure 9.**
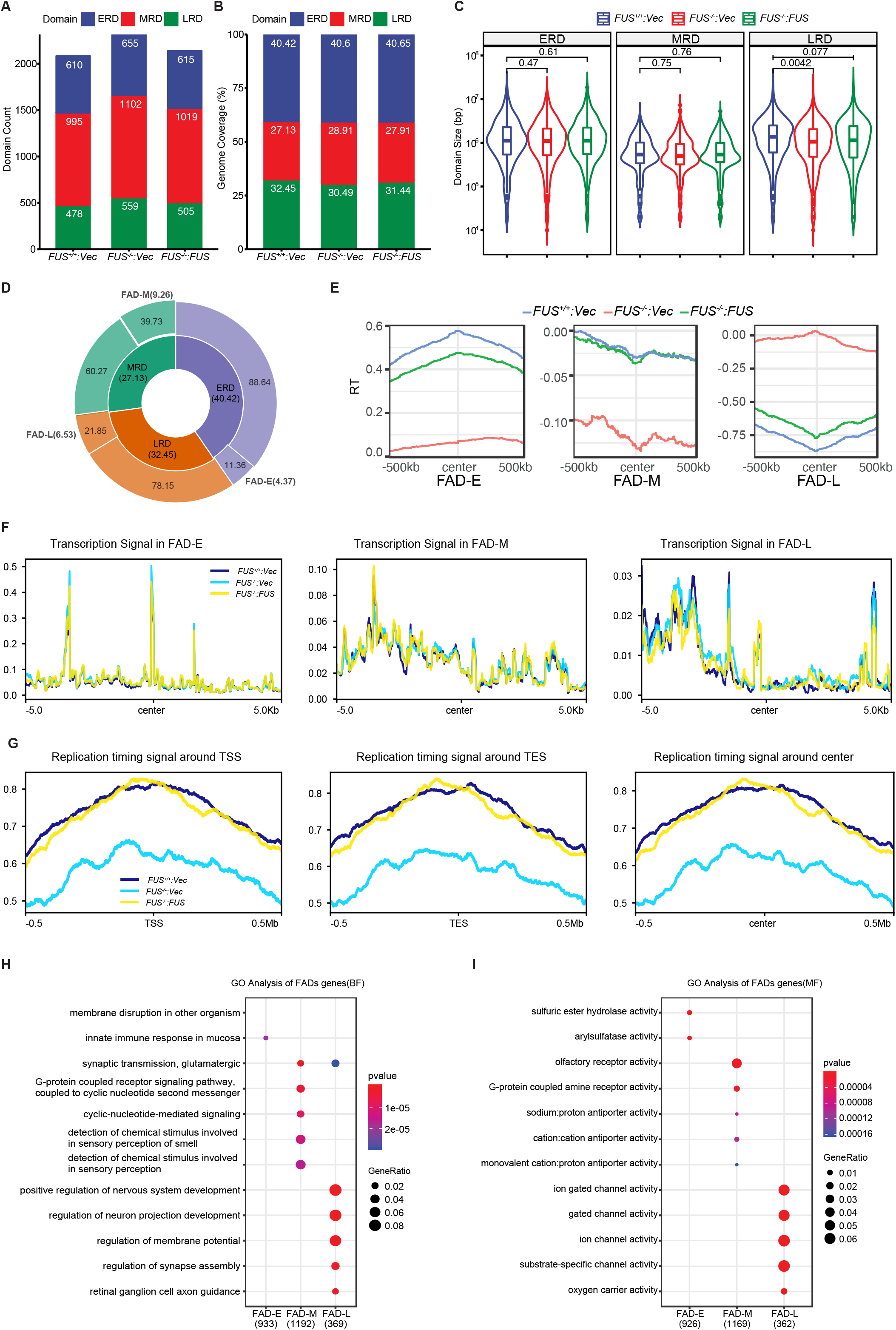
Characterization of FUS associated domains (FADs). A, RT profiles were segmented into three states by non-supervised package Segway as Early replication domain (ERD), Middle replication domain (MRD) and Late replication domain (LRD). The domain numbers in each sample were plotted and labeled. The two biological replicates were merged for replication domain segmentation. B, Percentages of genome coverage of replication domains in each sample were calculated based on the segmentation. The values are percentages of each domain. C, The same RT domain sizes are compared among all the samples. The Student *T*-test was used for determination of significance. D, Doughnut Pie chart of FAD coverage. The percentage of each RD (ERD, MRD, LRD; center pie) that is altered by FUS deficiency (FAD) is shown in the outside layer and the total percentage of each FAD (FAD-E, FAD-M, FAD-L) calculated and shown in parentheses. The percentage was calculated based on the genome coverage. E, RT signal enrichment analysis of FADs in the samples. The average domain size is ~10^6^ bp (C) and ~0.5 x 10^6^ bp flanking the midpoint was used for signal enrichment. Heatmap results of RT signal enrichment of changing ERD, MRD and LRD in all individual samples was shown in Sup. Fig. 8E to G. F, Transcription signal in the centered FADs. Transcription signal was normalized with CPM by STAR. G, RT signal enrichment around TSS, TES and center of FUS regulated gene regions across a ± 0.5Mb window. TSS: Transcription Start Sites; TES: Transcription End Sites. RT signal was calculated by log2 Ratio of S/G_1_ samples in 20kb bin after CPM normalization and followed with Z-score normalization. Only FUS regulated genes(listed in Table 4.) annotation was used. H, Gene ontology (GO) enrichment in biological function level of FADs. The FADs were extended 3000 bases in both ends and then the gene list under the extended FADs were extracted and was used for GO analysis. I, GO analysis in molecular function level of extended FADs.

To further characterize FADs, overlapping RDs in *FUS^+/+^* and *FUS^-/-^*:*FUS* cells were intersected and then subtracted from corresponding RDs in *FUS^-/-^* cells using bedtools. The resulting FADs (ERD-FUS, MRD-FUS, and LRD-FUS) comprised 11.36%, 39.73% and 21.85% of the total ERDs, MRDs, and LRDs in *FUS^-/-^* cells and represented 4.37%, 9.26% and 6.53% of whole genome sequence, respectively (Fig. 9D). In total, FADs covered ~20% of the genome in U-2OS cells. RT signals of FADs were centered and the distribution and heatmap analysis were performed and showed they were correctly identified (Fig. 9E and Sup. Fig. 8E to G).

Consistent with earlier studies (Hiratani et al., 2010; Hiratani et al., 2008; Rivera-Mulia et al., 2015), a positive correlation between gene activation and RT was found, with ERDs exhibiting active gene expression and LRDs exhibiting repressed gene expression (Fig. 9F and Sup. Fig. 8H). However, overall transcription signals were comparable between *FUS^-/-^* cells and *FUS^+/+^*cells across all three types of FADs (Fig. 9F). We next explored whether RT was changed proximal to genes showing FUS-dependent regulation by RNA-Seq. We found RT of annotated gene regions was delayed in *FUS^-/-^* cells relative to *FUS^+/+^* and *FUS^-/-^*:*FUS* cells (Fig. 9G). This pattern of delayed timing was observed across the entirety of the gene, including the transcription start site (TSS) and termination site (TES), and was observed for both upregulated and downregulated genes. To determine whether delayed RT was restricted to those genes regulated by FUS, we compared relative RT across all annotated genes. As shown in Sup. Fig. 8I and J, a similar pattern of delayed RT was observed in *FUS^-/-^* cells relative to *FUS^+/+^* and *FUS^-/-^*:*FUS* cells. These results imply that FUS plays a particularly important role in the early replication of transcriptionally active chromatin. Finally, we examined FADs for gene functional enrichment. Surprisingly given the non-neuronal nature of U-2 OS cells, we found LRD-FUS were highly enriched in nervous system development-related genes and, more specifically, genes encoding ion gated channels (Fig. 9H and I). These findings may be relevant to chromatin-associated functions of FUS in neurons.

### FUS is enriched in detergent insoluble nuclear structures

Given its role in RT, we wished to investigate the subnuclear localization of FUS in asynchronous cells and during S phase. Following detergent preextraction, endogenous FUS formed irregular nuclear puncta that in some instances were localized at the nuclear envelope. The FUS immunostaining signal was greatly reduced, but not absent, in *FUS^-/-^* cells, which may be due to limited cross-reactivity with EWSR1 or TAF15 (Fig. 10A). Stably expressed HA-FUS also localized to nuclear puncta (Sup. Fig. 9A). However, FUS did not colocalize with RIF1, a master regulator of RT (Sup. Fig. 9B). The latter finding suggests that FUS and RIF1 regulate RT through distinct mechanisms. FUS puncta were not diminished by RNAase treatment but were largely abolished by the RNA/DNA nuclease Benzonase (Fig. 10B). To investigate whether endogenous FUS puncta represent phase-separated liquid droplets reported in other studies (Altmeyer et al., 2015; Chong et al., 2018; Kroschwald et al., 2015; Kroschwald et al., 2017; Li et al., 2013; Patel et al., 2015; Peskett et al., 2018), we treated U-2 OS cells with 1,6-hexanediol(1,6-HD). 1,6-HD failed to disperse FUS puncta even after long incubation periods, suggesting that these structures do not meet the operational definition of liquid droplets (Fig. 10C).

**Figure 10.**
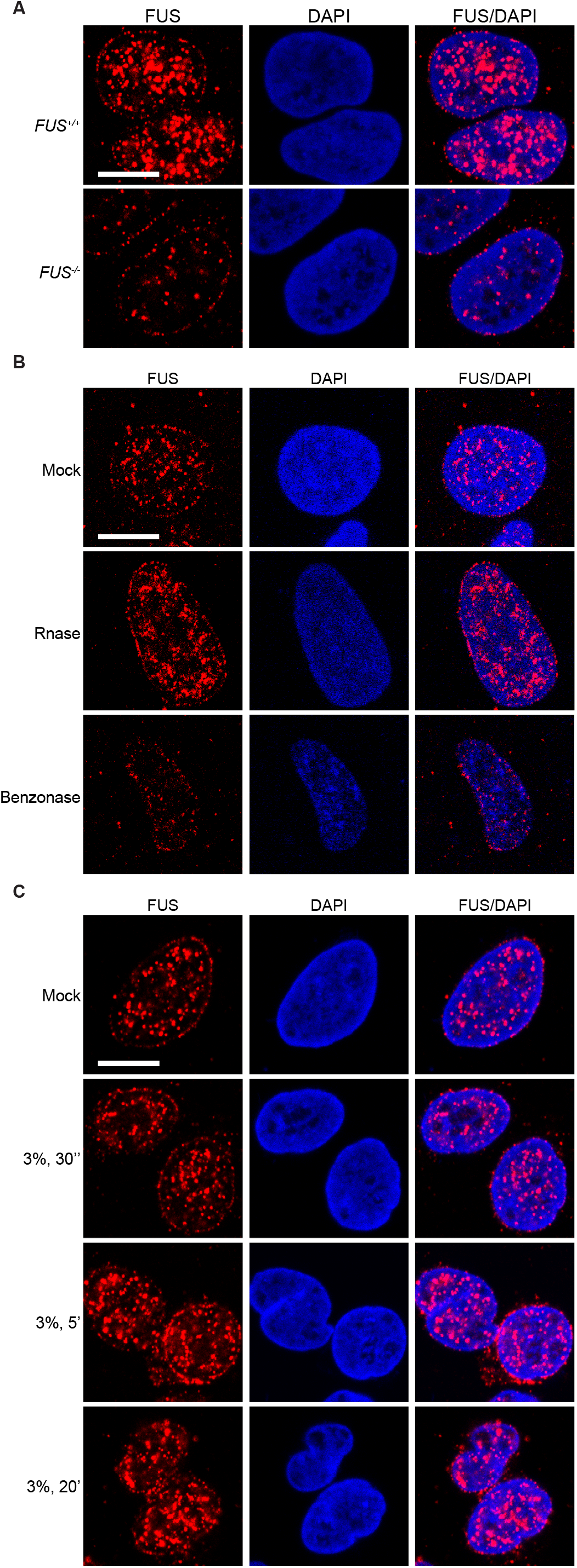
RNA-binding independent association of FUS with chromatin. A, *FUS^+/+^* and *FUS^-/-^* cells were pre-extracted with CSK buffer, fixed, and stained with α-FUS. B, U-2 OS cells were pre-extracted in CSK buffer containing 0.3mg/ml Rnase A or 100U/ml Benzonase and stained with α-FUS. C, U-2 OS Cells were treated with 3% 1,6-HD for indicated times and then processed for immunofluorescence staining with FUS antibodies.

## Discussion

FUS DNA repair functions have been deduced from its PARP-dependent recruitment to sites of microirradiation; its coimmunoprecipitation with repair proteins; and the modest chromosome instability and DSB repair defects of FUS-deficient cells (Deng et al., 2014; Hicks et al., 2000; Martinez-Macias et al., 2019; Mastrocola et al., 2013; Rulten et al., 2014; Singatulina et al., 2019; Wang et al., 2018; Wang et al., 2013). Despite these results, a unifying role for FUS in genome protection has yet to emerge. Using *FUS^-/-^* cells with and without reconstitution we found that, while FUS plays a supporting role in DSB repair, it is prominently involved in both the initiation and timing of DNA replication. DNA replication defects plausibly contribute to genome instability and DDR-related phenotypes ascribed to FUS-deficient cells.

FUS is among the first factors recruited to sites of microirradiation, which is driven through association of FUS RGG domains with PAR chains (Altmeyer et al., 2015; Mastrocola et al., 2013; Rulten et al., 2014; Singatulina et al., 2019; Wang et al., 2013). FUS is also reportedly required for the assembly of IR-induced 53BP1 foci (Wang et al., 2013), despite the fact that FUS does not accumulate at these structures (Mastrocola et al., 2013). Results in Sup. Fig. 1A-B clearly show that γH2AX and 53BP1 recruitment occurs independently of FUS in U-2 OS cells, with similar findings made for *FUS^-/-^* H460 cells (not shown). In fact, 53BP1 foci were significantly larger and more persistent in *FUS^-/-^* cells relative to *FUS^+/+^* controls (Sup. Fig. 1A). This result appears to be congruent with findings of Altmeyer et al. who reported that overexpression of the EWSR LCD suppressed IR-induced 53BP1 focus formation (Altmeyer et al., 2015). It is conceivable that FUS inhibits local assembly of 53BP1 complexes and/or limits their lateral spread along damaged chromatin. Non-exclusively, persistent 53BP1 foci may reflect delayed kinetics of DSB repair in *FUS^-/-^* cells. Chromatin-bound FUS notably interacted with DNA-PK and its Ku70 and Ku80 DNA-binding subunits that mediate NHEJ; however, *FUS^-/-^* U-2 OS cells were not appreciably radiosensitive, indicating FUS is not an essential NHEJ factor.

Consistent with their slow growth phenotype, *FUS^-/-^* cells exhibited reduced RF speed (Fig. 4B), delayed RF restart (Fig. 4C), reduced expression of S-phase-associated genes (Fig. 3C), and reduced loading of pre-RC complexes (Fig. 5B). A potentially direct role for FUS in DNA replication was suggested by the presence of DNA replication factors in FUS-chromatin complexes (Fig. 6). The association of FUS with lagging strand synthesis factors POLδ1, PCNA and FEN1, but not leading strand POLε, further suggested that FUS may play a role in the deposition or removal of RNA primers and/or the ligation of single-strand nicks on the lagging strand. It is worth noting that PARP, which was also present in FUS-chromatin complexes, contributes to the ligation of Okazaki fragments on the lagging strand (Hanzlikova et al., 2018). Despite these interactions, FUS did not stably associate with translocating replisomes in the iPOND assay (Fig. 4D). Although this does not necessarily rule out a direct role for FUS at RFs, we speculate that FUS impacts DNA replication are a reflection of its impacts on local chromatin structure and transcription.

To our knowledge, this is the first study to implicate FUS in the control of RT. The RT program is a stable, cellular characteristic that is established in early G_1_ phase(Dileep et al., 2015; Dimitrova and Berezney, 2002; Dimitrova and Gilbert, 1999). Spatiotemporal control of RT is highly dependent on the master timing factor RIF1, a chromatin-bound factor that also plays important roles in RF stabilization and DSBR pathway choice (Buonomo, 2017; Dev et al., 2018; Escribano-Diaz et al., 2013; Gupta et al., 2018; Mirman et al., 2018; Noordermeer et al., 2018). RIF1-deficient mammalian cells or yeast exhibit spatial changes in DNA replication that correlate with premature replication origin firing (Cornacchia et al., 2012; Hayano et al., 2012;

Silverman et al., 2004; Xu et al., 2010; Yamazaki et al., 2012). Recent studies have correlated genome-wide RIF1 chromatin occupancy with the RT of individual chromatin domains and established a role for RIF1 in the bundling of coregulated origins, termed RIF1-associated domains, or RADs (Foti et al., 2016). Beyond RIF1, few genetic regulators of RT have been identified(Fernandez-Vidal et al., 2014). Our findings suggest that FUS acts bidirectionally to control RT in a chromosome context-dependent manner. The fact that some chromosomal domains replicate earlier in *FUS^-/-^* cells relative to *FUS^+/+^* cells suggests that the RT functions of FUS are at least partially independent from its positive contributions to DNA replication initiation.

Two plausible models may underlie participation of FUS in RT, with both models invoking the phase-separation characteristics of FUS (Boeynaems et al., 2018) as a central mechanistic feature. First, FUS may fulfill a chromatin-bundling function (Foti et al., 2016; Yamazaki et al., 2012). In this model the DNA binding and dynamic oligomerization properties of FUS promote the assembly of FADs that are replicated with similar timing. This role would be conceptually similar to that proposed for RIF1, though the distinct localization profiles of FUS and RIF1 strongly suggests they regulate RT through different pathways.

Non-exclusively, we favor a model in which FUS-dependent RT is linked to its roles in transcriptional activation. The LCDs of FUS, EWSR1, and TAF15 bind to the CTD of RNA Pol II (Bertolotti et al., 1998; Burke et al., 2015; Kwon et al., 2013; Schwartz et al., 2012) and function as potent transcriptional activators when fused to heterologous DNA-binding domains(Bertolotti et al., 1999; May et al., 1993; Zinszner et al., 1994). Indeed, transcriptional deregulation is thought to drive malignant transformation in soft-tissue sarcomas harboring oncogenic fusions of FET genes with site-specific transcription factors such as CHOP, FLI1, and CREB(Antonescu et al., 2006; Bailly et al., 1994; Crozat et al., 1993). A reciprocal relationship between RT and transcription is supported by studies showing that RT switches during embryonic development precede transcriptional changes of proximal genes (Kaaij et al., 2018; Rivera-Mulia et al., 2015; Siefert et al., 2017) and work showing that transcriptional activation leads to RT advancement (Therizols et al., 2014). Sima et al. further demonstrated that *cis*-regulatory elements within an enhancer promoted early RT of the *Dppa2/4* domain in mouse ESCs (Sima et al., 2018). While absolute levels of transcription were not significantly different, transcriptionally-active genes showed delayed RT in *FUS^-/-^* cells relative to *FUS^+/+^* cells (Fig. 9F and G). The LCD of FUS, in addition to intrinsically disordered regions of transcriptional coactivators BRD4 and MED1 have been implicated in the assembly of phase-separated transcription “condensates” at gene enhancers (Cho et al., 2018; Chong et al., 2018; Sabari et al., 2018). We speculate that FUS contributes to clustering of transcription complexes and chromatin looping that ultimately specifies FADs that undergo coordinate RT regulation. Chromatin capture approaches, such as ChIA-PET, will be needed to test this hypothesis and to investigate the potential role for PAR-dependent FUS liquid demixing in RT and DNA replicaion control (Altmeyer et al., 2015; Singatulina et al., 2019). In conclusion, our findings establish new roles for FUS in whole genome wide DNA replication and timing regulation that may contribute to broader genome instability deficits of FUS-deficient cells.

## Materials and Methods

### Cell culture and gene editing

The U-2 OS, H460 and HEK293T cell line was obtained from the American Type Culture Collection (ATCC). U-2 OS and U-2 OS derivative cell lines were grown in McCoy’s medium (Corning, 10-050-CV). H460 cells were grown in RPMI-1640 medium (Corning, 10-043-CV). HEK293T cells were grown in DMEM medium (Corning, 10-013-CV). All cell lines were grown in its medium with 10% fetal bovine serum (Atlanta biologicals) and 1% Penicillin/Streptomycin (Corning, 30-002-CI) and incubated at 37°C in 5% CO_2_. For G_1_/S boundary via double thymidine block. Briefly, cells were treated with 2mM thymidine for 19 hours and 16 hours with a 9 hours interval of growth without thymidine. Cells were washed three times with PBS, and then released into S phase and harvested at indicated time points.

*FUS^-/-^* cells were generated by transient transfection of U-2 OS cells with pX459 (v2, Addgene plasmid #62988) vectors(Ran et al., 2013) expressing sgRNAs (CGCCAGTCGAGCCATATCCC and AGAGCTCCCAATCGTCTTAC) targeting exon 4 using jetPRIME (Polyplus). Twenty-four hours after transfection, cells were selected for 72h with 1μg/ml puromycin and then cells were diluted to 96-well-plate with 1cell/well and single clones were isolated and screened for FUS knockout by Western blotting. All clones were sequenced around the targeted sequence and four clones were selected for further study. We reconstituted *FUS^-/-^* Cl.110 with a FUS CDS cloned into a pQCXIH CMV/TO DEST retroviral vector (addgene, #17394) vector by Gateway cloning. As a negative control, the GUS gene from vector pENTRGUS (invitrogen) was also cloned into pQCXIH CMV/TO DEST vector. Retroviral plasmids were packaged with GP2-293 packaging cell line (Clonetech, 631458). Stably transduced cells were selected with 50μg /ml hygromycin for 1 week and single clones isolated, expanded, and tested for FUS expression.

### EdU labeling, flow cytometry, microscopy, and DNA fiber analysis

For cell cycle progression experiments U-2 OS cells were incubated with 20 μM EdU for 30 min before collection and then fixed with ice-cold 70% ethanol. EdU detection was performed using the Click-IT Plus EdU Alexa Fluor 647 Flow Cytometry Assay Kit (Life Technologies, C10634). Propidium (PI) was added to a concentration of 50 μg/ml. Flow cytometry was performed on Thermo Fisher Attune, and data was analyzed and organized using FlowJo software. For in situ EdU and BrdU staining, U-2 OS cells were pulse labelled for 30 min with 20μM BrdU or EdU and fixed with 4% paraformaldehyde (PFA). For BrdU detection, cells were then incubated with 2M HCl for 30 min and then permeabilized with 0.2% Triton-X100 for 15 min at room temperature, washed and blocked in 3% BSA. Cells were stained with BrdU primary antibody (Santa Cruz, sc-32323) in 3% BSA and incubated overnight in 4°C, followed by washing in PBST (PBS with 0.02% Tween-20) and incubation with appropriate secondary antibodies in 3% BSA for 1 h at room temperature. EdU detected by click chemistry and described above. Samples were mounted in VECTASHIELD mounting medium with DAPI (Vector, H-1200) before imaging. For general immunostaining experiments, cells were seeded into 12-well-plate with glass coverslip (and transferred to a humidity chamber for immunostaining with appropriate antibodies. Nuclear DNA either stained with 0.5μg/ml DAPI for 10 min at room temperature and then mounted with mounting medium for fluorescence (Vector, H-1000), or directly mounted in mounting medium with DAPI for fluorescence (Vector, H-1200) before imaging. Images were acquired using a Nikon A1RS Confocal Microscope under a 63x oil immersion objective. Images were organized using Fiji ImageJ software. PLA foci were counted in CellProfiler (version 3.1.5). DNA fibers were prepared and analyzed as described in (Tonzi et al., 2018). In brief, cells were pulsed with 50 μM IdU and CldU for times indicated in each experiment. Cells were lysed directly on glass slides, fixed, denatured, stained, and imaged with Keyence BZ-X710 microscope. Image analysis were done with ImageJ. A minimum of 150 fibers were measured for each independent experiment and analysis shows mean of three independent experiments (biological replicates).

### RNA-Seq and gene expression

Total RNA was isolated using the TRIzol reagent (Invitrogen, 15596018) following the manufacturer’s protocol and treated with TURBO Dnase (Invitrogen, AM2239). Then RNA samples were sent to Novogene (Novogene Co., Ltd, Sacramento, CA) for non-stranded cDNA library building and sequencing at PE150 with NovoSeq 6000. Raw reads adapters were trimmed by fastp(Chen et al., 2018b) and then were mapped to human genome(GRCh38) by STAR with the setting suggested by ENCODE project (https://github.com/ENCODE-DCC/rna-seq-pipeline). The number of RNA-Seq reads mapped to each transcript were summarized with featureCounts(Liao et al., 2014) and differential expression was called using DESeq2(Love et al., 2014). Three biological replicates were used for each sample. For qPCR analysis total RNA was reverse transcribed into cDNA using SuperScript IV VILO Master Mix with ezDnase enzyme Kit (Invitrogen, 11766050). The primers were designed by Beacon Designer or NCBI primer-blast online tool. q-PCR reaction was performed on Bio-Rad CFX RealTime PCR system using iTaq Universal SYBR Green Supermix (Bio-Rad, 1725125)

### Replication timing analysis

Cells were prepared and collected accordingly to (Koren et al., 2012) with the following modifications. Around 10 million asynchronous cells were collected and fixed in 70% ethanol. Fixed cells were washed with ice-cold PBS and treated with Accutase (CORNING, 25-058-CI) for 20 minutes at room temperature. Cells were pelleted and resuspended in 2 ml PBS with 250 μl 10 mg/ml RNaseA and incubated at 37°C for 30 minutes and stained with propidium iodide (PI), and then sorted to G_1_ and S-phase fractions by flow cytometry. DNA extracts from sorted cells were prepared using with DNeasy Blood and Tissue Kit (Qiagen, 69504) and single-end 100-base sequencing libraries prepared using TruSeq kit (Illumina) and deep sequencing was performed on HighSeq 2500. The analysis was carried out according to Marchal et al. (Marchal et al., 2018). Briefly, Reads were trimmed by fastp and then were mapped onto the human genome (GRCh38) using bowtie2. The replication timing (RT, S/G_1_ ratio) was calculated in a fixed window size of 20Kb. Then RT raw data were used for quantile normalization, and then smoothened with Loess smoothing. The RT signal and replication signal enrichment analysis were performed by deeptools(Ramirez et al., 2016).Two biological replicates were analyzed separately.

### Immunoblotting

For whole-cell extraction, cells were resuspended in high salt lysis buffer (50 mM Tris, pH 7.5, 300 mM NaCl, 10% glycerol, 0.5% Triton X-100, 2mM MgCl_2_, 3mM EDTA, 1% Protease Inhibitor Cocktail (Sigma, P8340-5ml) supplemented with Benzonase (50 U/ml) and incubated on ice for 20 min followed by the addition of 4X SDS-loading buffer and heating at 95°C for 15 min. For chromatin fractionation, cells were resuspended in cytoskeleton (CSK) buffer (20 mM HEPES-KOH (pH 7.4), 100 mM NaCl, 3 mM MgCl_2_, 300 mM sucrose and 1% Protease Inhibitor Cocktail(Sigma, P8340-5ml)) containing 0.5% Triton X-100, incubated on ice for 20 min, and centrifuged for 5min at 5,000 x g at 4 °C. The supernatant was transferred to a new tube and saved as soluble fraction (SF), while the pellet/chromatin fraction, (CF) was washed twice in CSK buffer without detergent and resuspended in CSK buffer with Benzonase (50U/ml) for 20 min digestion at which time 4 X SDS loading buffer was added and the lysates heated to 95°C for 15 min. For immunoblotting, samples were separated by SDS-PAGE and transferred to PVDF membranes and immunoblotted with primary antibodies and LI-COR IRDye secondary antibodies (IRDye 800CW goat anti-rabbit and IRDye 680RD goat anti-mouse) as described (Kim et al., 2018; Kim et al., 2016). Signals were acquired using Odyssey bio-systems (LI-COR Biosciences). Immunoblotting results were analyzed and organized with ImageStudio Lite software (LI-COR).

### FUS purification and mass spectrometry

RIME (rapid immunoprecipitation mass spectrometry of endogenous proteins) assay of FUS were carried out as described (Mohammed et al., 2016) with the following modifications. Briefly, around 20 million cells were counted and fixed with 20 ml 1% formaldehyde solution for 8 minutes at room temperature. Fixation was quenched by adding 0.12M Glycine. The soluble fraction was extracted in 10ml of LB1 (50 mM HEPES-KOH (pH 7.5), 140 mM NaCl, 1 mM EDTA, 10% Glycerol, 0.5% NP-40, 0.25% Triton X-100, 1% Protease Inhibitor Cocktail (Sigma, P8340-5ml)) for 10 min with rotation at 4°C. Cell nuclei were pelleted and washed once with 10ml LB2 (10 mM Tris-HCl(pH 8.0), 100 mM NaCl, 1 mM EDTA, 0.5 mM EGTA, 1% Protease Inhibitor Cocktail) and then resuspended in 500 μl LB3 (10 mM Tris-HCl (pH 8.0), 100 mM NaCl, 2.5 mM MgCl_2_, 0.1% (W/V) sodium deoxycholate, 0.5% Triton X-100, 1% Protease Inhibitor Cocktail) with 500 U Benzonase and incubated at room temperature for 30 min. Benzonase was deactivated with 2 mM EDTA, 1 mM EGTA. To this mixture was added 50 μl 10% Triton X-100, 37.5 μl of 4 M NaCl, and LB3 to bring the total lysate volume of each sample to 1 ml. Digested lysates were briefly sonicated using a 10s/50s on/off cycle for three times at 40% power and clarified by centrifugation at 20,000g for 10 min at 4°C and supernatants were incubated with 10 μg FUS antibody (Bethyl, A300-302A) overnight at 4°C with rotation. Subsequently, 50 μl of pre-washed Dynabeads protein G (Invitrogen, 10003D) was added to the lysates and incubated for additional 4 h at 4°C. For western blot, beads were washed sequentially with 1 ml LB3 and 1 ml RIPA buffer (50 mM HEPES-KOH (pH 7.5), 0.5 M LiCl, 1 mM EDTA, 1% NP-40, 0.7% (W/V) sodium deoxycholate, 1% Protease Inhibitor Cocktail) once and boiled in 100μl 2 x SDS buffer. For mass spectrometry, beads were washed 5 times with 1 ml RIPA buffer and twice in 1 ml of cold fresh prepared 100 mM ammonium hydrogen carbonate (AMBIC) solution and processed as described (Mohammed et al., 2016).

FUS RIME IPs from *FUS^+/+^* and *FUS^-/-^* cells were subjected to tryptic digestion and orbitrap mass spectrometry (MS) using the filter aided sample preparation (FASP) method (Wisniewski et al., 2009). We performed two technical replicates for each of the three biological replicates. The MetaMorpheus software program was used to identify peptides and proteins in the samples(Shortreed et al., 2015; Solntsev et al., 2018). Protein fold changes were quantified by FlashLFQ(Millikin et al., 2020; Millikin et al., 2018).

### Cell proliferation, survival, and iPOND assays

For cell proliferation assay, 500 cells were plated in each well of 96-well-plate and each sample had 6 replicates and monitored for 6 days from day 0 to day 5 by CellTiter-Glo 2.0 Assay (Promega, G9242) according to the manufacturer’s instructions. The luminescence was recorded by SpectraMax i3 (Molecular Devices). For cell viability assay, 1000 cells/well were plated in 96-well-plate with drug-free medium and varying amounts of drugs were added after 12 h in fresh medium. Cell survival was assayed as same as cell proliferation assay after 3 or 5 days as indicated in figure legend. Data was analyzed and organized by Prism 7.

### iPOND assay

The iPOND experiments were performed as described in (Dungrawala and Cortez, 2015; Tonzi et al., 2018) with minor modifications. Briefly, ~10^8^ cells were pulse-labelled with 20 μM EdU for 15 min followed by a 1 h chase with 20 μM thymidine. To induce replication stress, cells were treated with 2 mM Hydroxyurea (HU) after EdU labeling for 2 h, and then chased with 20 μM thymidine for 1 h. Each plate was crosslinked with 10 ml 1% formaldehyde in PBS for 20 min and quenched by adding 1ml of 1.25 M glycine for 5 min. The conjugation of biotin to EdU was carried out by click chemistry reaction for 2 hours at room temperature in click reaction buffer(10 μM biotin-azide, 10mM sodium-L-ascorbate, 2mM CuSO4 and 800μM THPTA in PBS) and followed by washing once in 0.5% BSA in PBS and once in PBS. Cells were resuspended in LB3 with 500U Benzonase (Santa Cruz, sc-202391) and incubated at room temperature for 30 minutes. Digested lysates were briefly sonicated using a 10s/50s on/off cycle for four times at 40% power and clarified by centrifugation at 8,000g for 10min at 4°C and supernatants were incubated overnight with 50μl magnetic streptavidin beads (Dynabeads MyOne Streptavidin T1, 65601) at 4°C with rotating. Beads were washed once in 1ml washing buffer (20mM Tris-HCl(pH 8.0), 500mM NaCl, 2mM EDTA, 0.1%(W/V) sodium deoxycholate,1% Triton X-100), once with 1ml RIPA buffer(50mM HEPES-KOH(pH 7.5), 0.5M LiCl, 1mM EDTA, 1% NP-40, 0.7%(W/V) sodium deoxycholate, 1% Protease Inhibitor Cocktail) and twice in LB3 buffer, and then proteins were eluted by boiling in 2 x SDS buffer for 25 min.

### Statistical processing

Statistical analysis information including individual replicates and biological replicates number, mean or median, and error bars are explained in the figure legends. The statistical tests and resulting *P* values are showed in the figure legends and/or figure panels.

**Key resources table.**
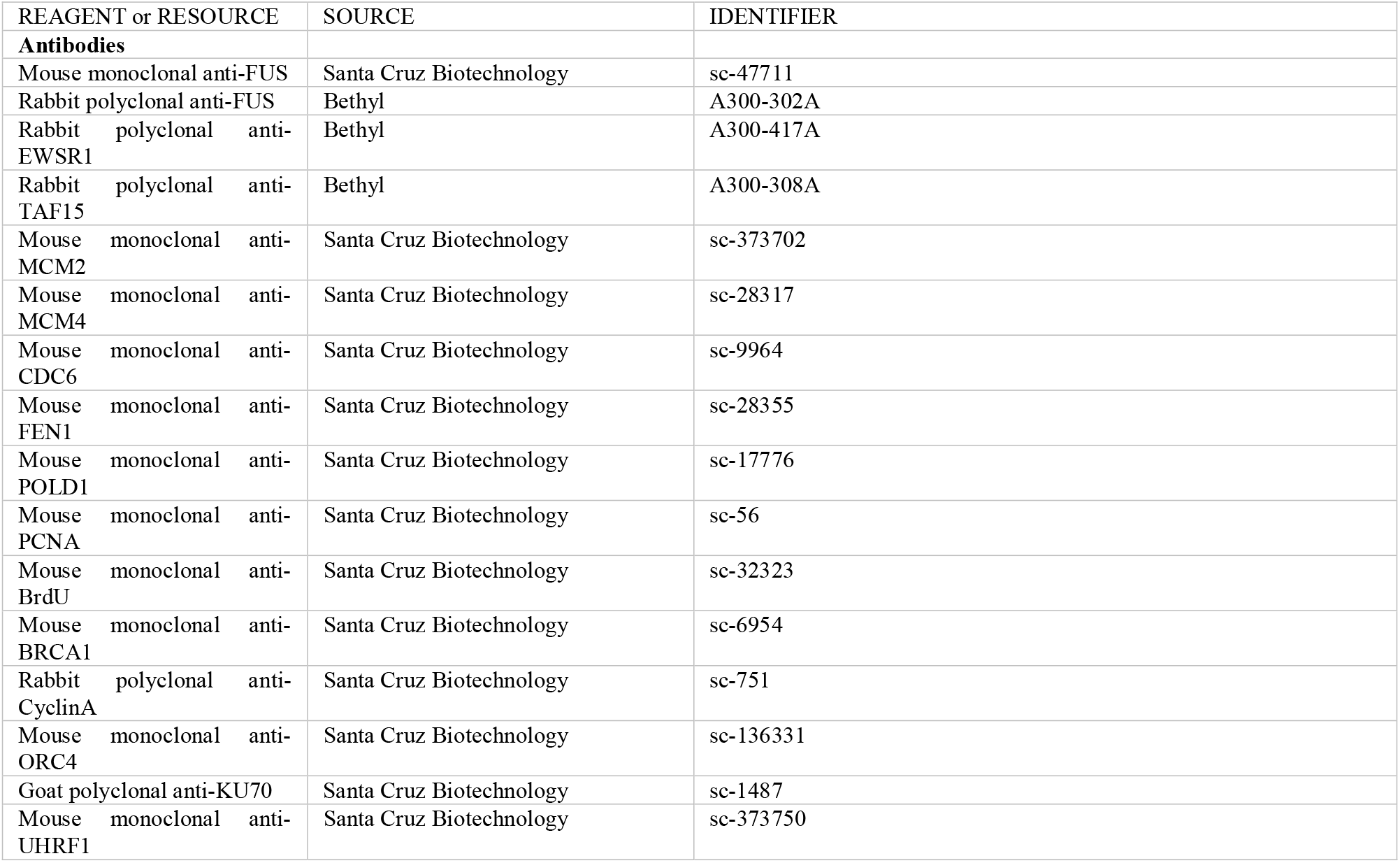
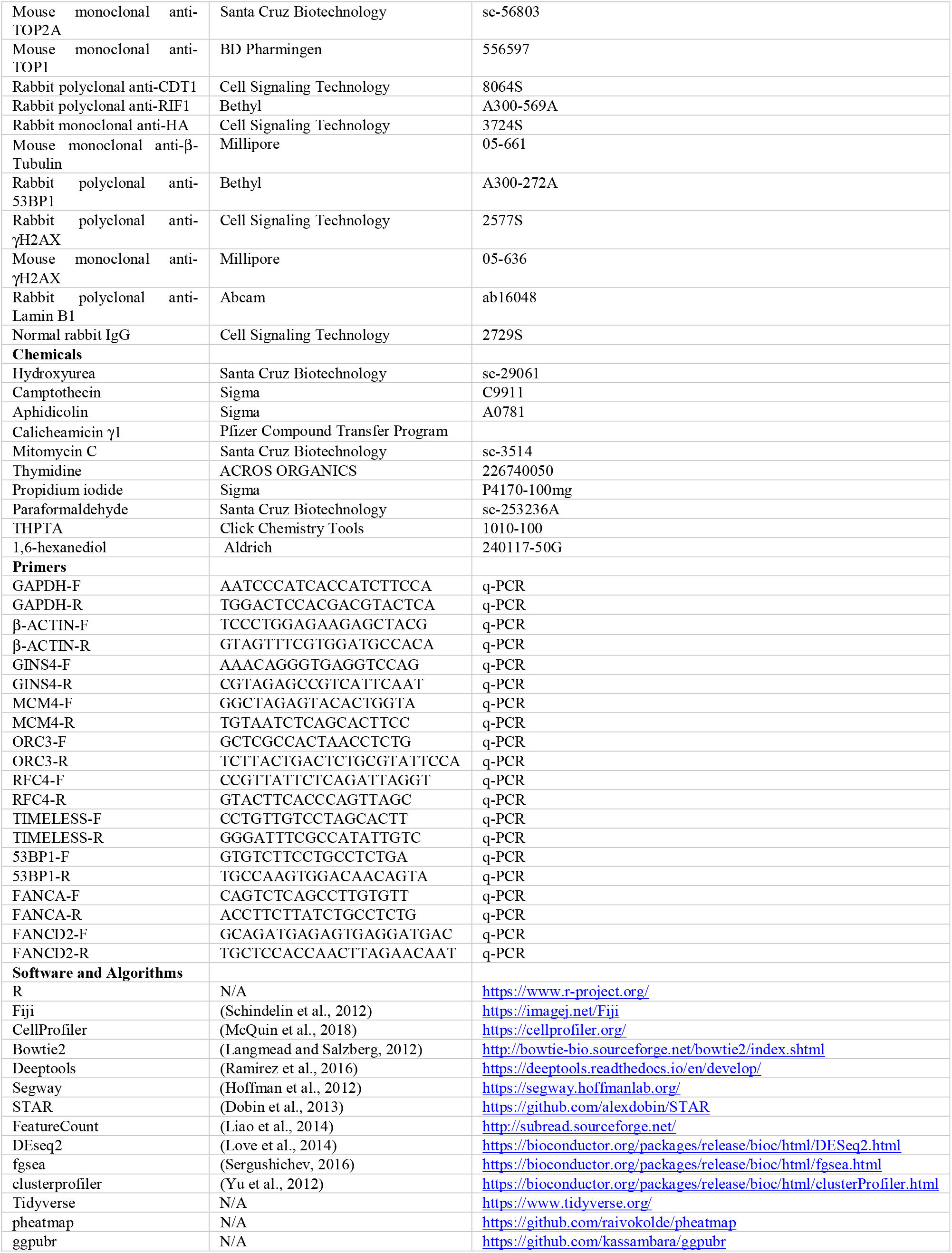

## Data availability

All the source data represented in the figures and bioinformatic analysis scripts are available on github (https://github.com/biofisherman/FusReplication). The accession number of Replication timing sequencing data in NCBI: PRJNA615974. The GEO number of RNA-Seq data: GSE147784.

## Acknowledgments

We gratefully thank Ammon Koren provided protocol for replication timing sample preparation and communication of replication timing analysis. We are grateful to Markku Varjosalo for pcDNA5/FRT/TO/StrepIII/HA/GW vector, Pfizer Compound Transfer Program for Calicheamicin A. We thank Lance A Rodenkirch in optical imaging core and Flow Cytometry Laboratory for assistance.

## Supplemental Figures

**Supplemental Figure 1.**
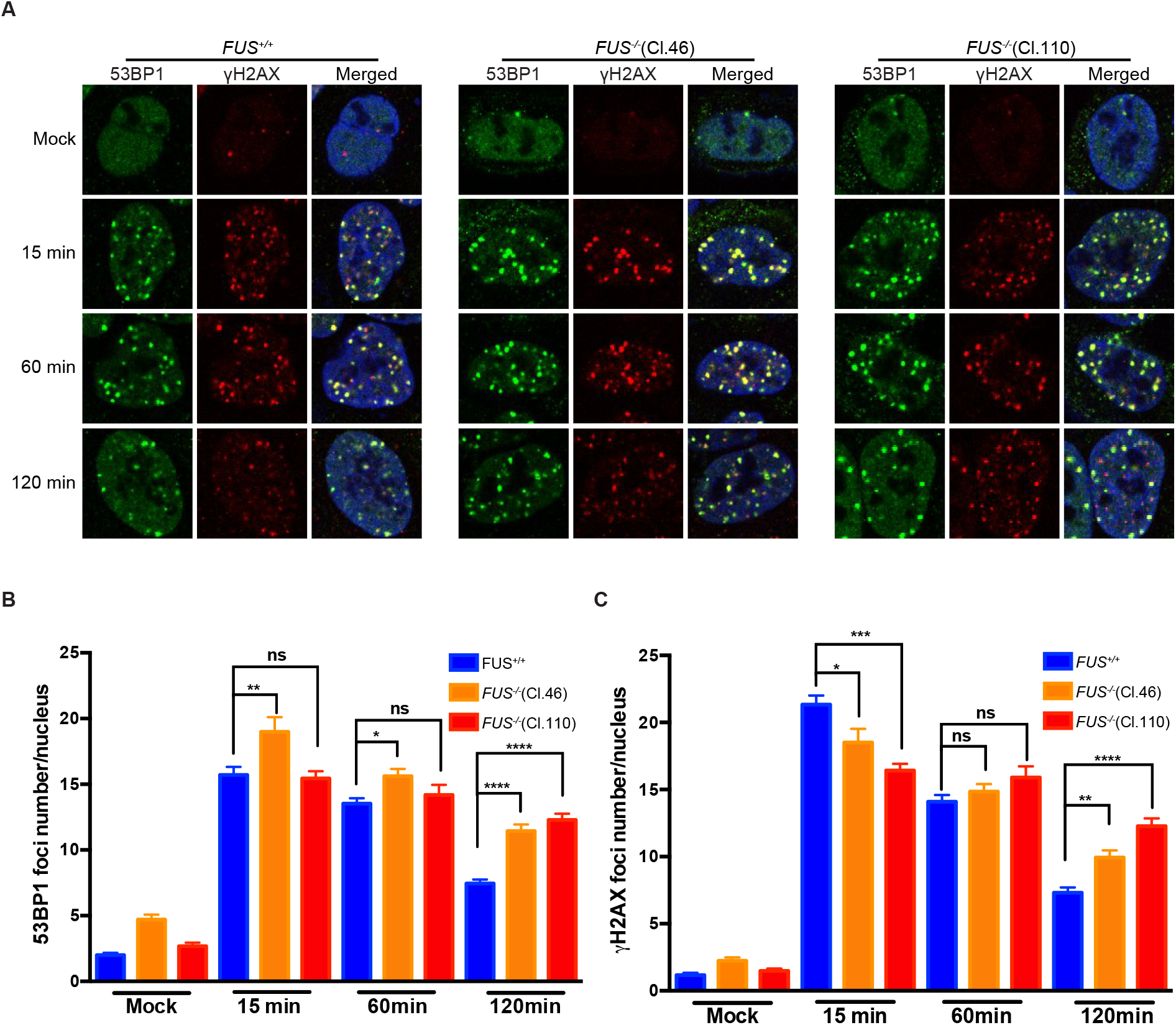
Increased persistence of 53BP1 and *γ*H2AX foci in irradiated *FUS^-/-^* U-2 OS cells. A, Representative 53BP1 and γH2AX staining patterns of *FUS^+/+^* cells and two *FUS^-/-^*, clones (Cl.46 and Cl.110), harvested at different times after exposure to 2 Gy IR.. *B* and *C*, Quantification of 53BP1 (B) and γH2AX (C) foci. At least 50 cells were analyzed for each sample. The bars represent mean with *SEM.* Significance was calculated using one-way ANOVA(\*\*\*\**P*<0.0001) followed by Bonferroni’s Multiple Comparison Test.

**Supplemental Figure 2.**
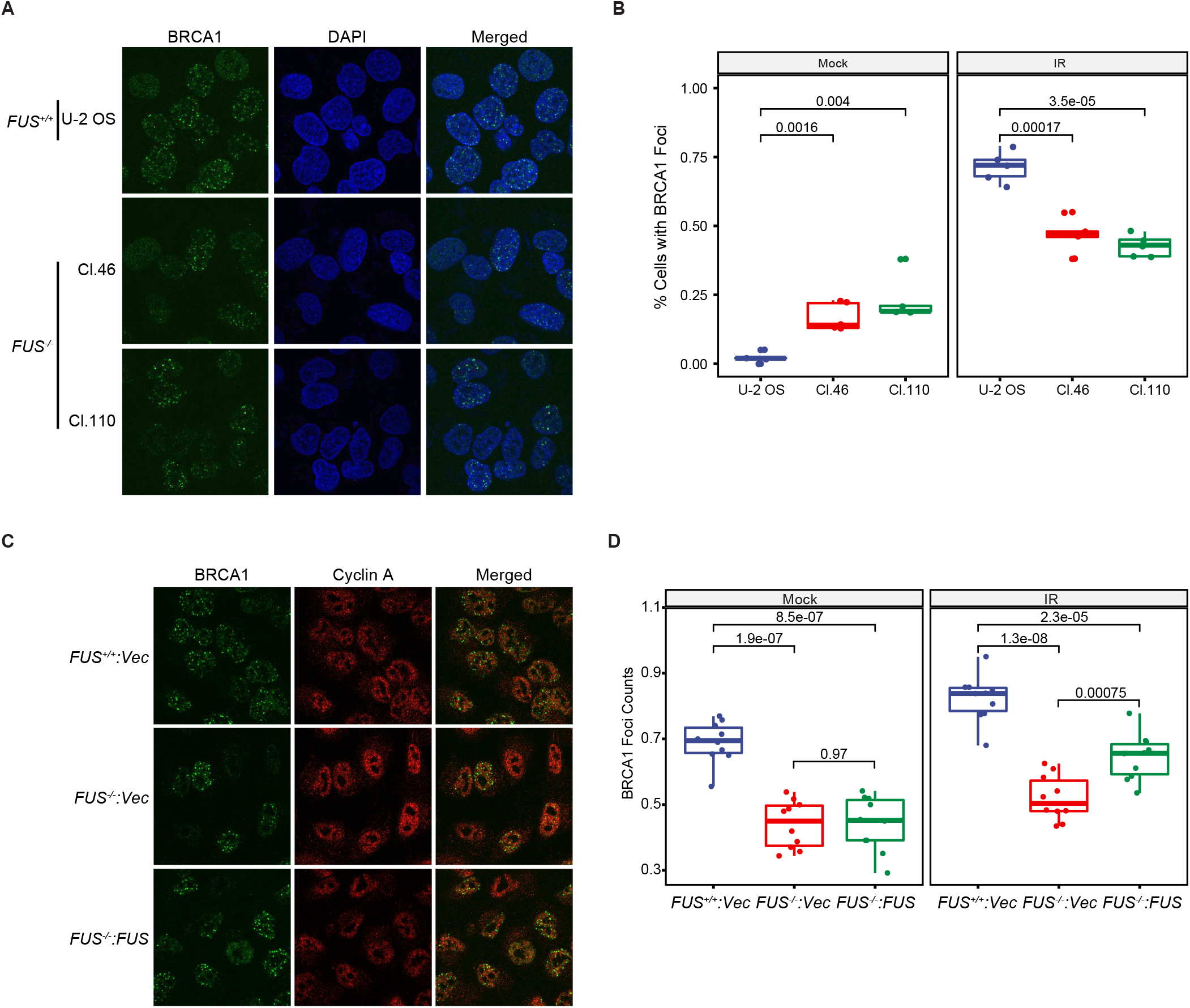
FUS promotes BRCA1 recruitment to DNA damage sites. A, *FUS^+/+^* and *FUS^-/-^* U-2 OS cells were harvested prior to or 15 min after exposure to 2 Gy IR and stained with BRCA1 antibodies. B, Cells displaying five or more BRCA1 foci were tabulated and subjected to statistical treatment using student *T-test* (n=5 five random regions). C, FUS reexpression reverses the BRCA1 focus formation defect *FUS^-/-^* cells in of S/G_2_-phase. Cells were fixed and stained with BRCA1 and Cyclin A2 antibodies before or 15 min after exposure to 2 Gy IR. D, Numbers of BRCA1 foci in cyclin A-positive cells were tabulated. Student *t-test*, n=10 (random regions).

**Supplemental Figure 3.**
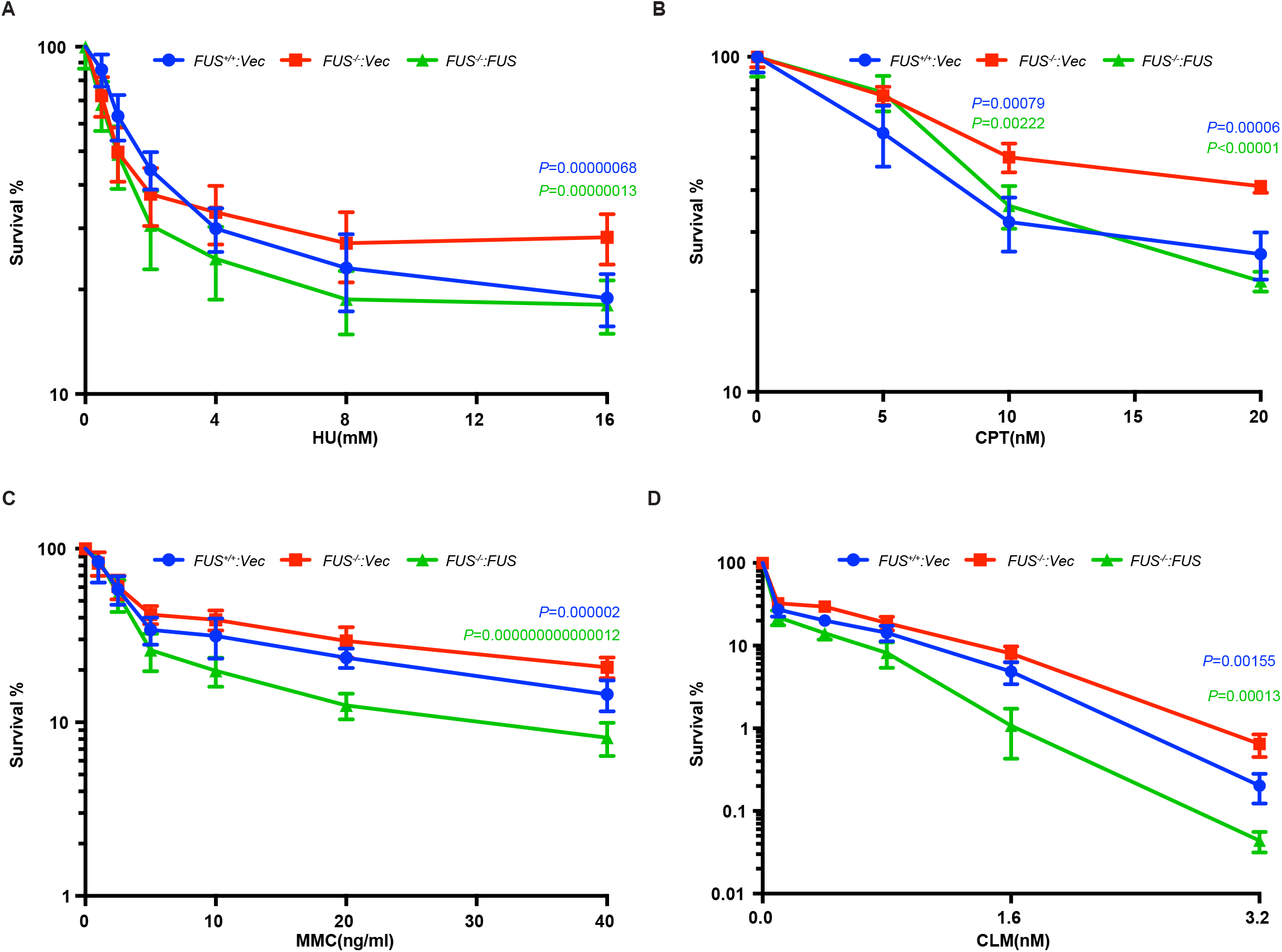
Sensitivity of *FUS^-/-^* cells to genotoxins. A to D, CellTiter-Glo assay was performed to measure the cell survival after 5-days continuous treatment with HU, CPT, MMC or CLM at the indicated concentration. 1000 cells were plated in each well. HU and MMC treatments employed three biological replicates of5 technical replicates each. CPT and CLM employed one biological replicate with 5 technical replicates. The bars represent mean ± SD. *P* values were calculated suing *t*-test. The blue *p*-values compare *FUS^+/+^*:*Vec* and *FUS^-/-^*:*Vec* and the green p-values compare *FUS^-/-^*:*FUS* to *FUS^-/-^*:*Vec*.

**Supplemental Figure 4.**
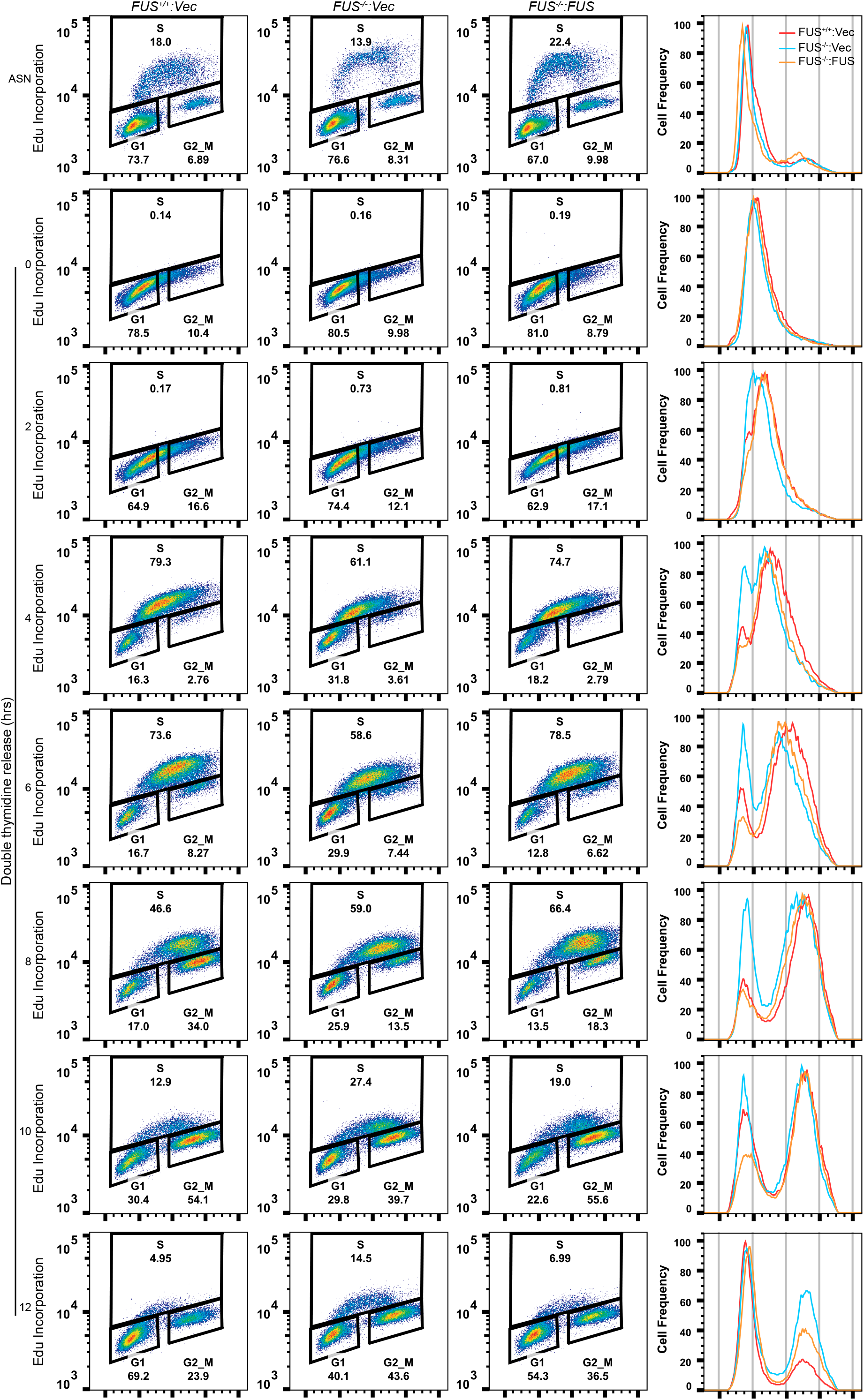
Full time course analysis of cell cycle progression in *FUS^+/+^*, *FUS^-/-^* and *FUS^-/-^*:*FUS* cells shown in Fig. 2B.

**Supplemental Figure 5.**
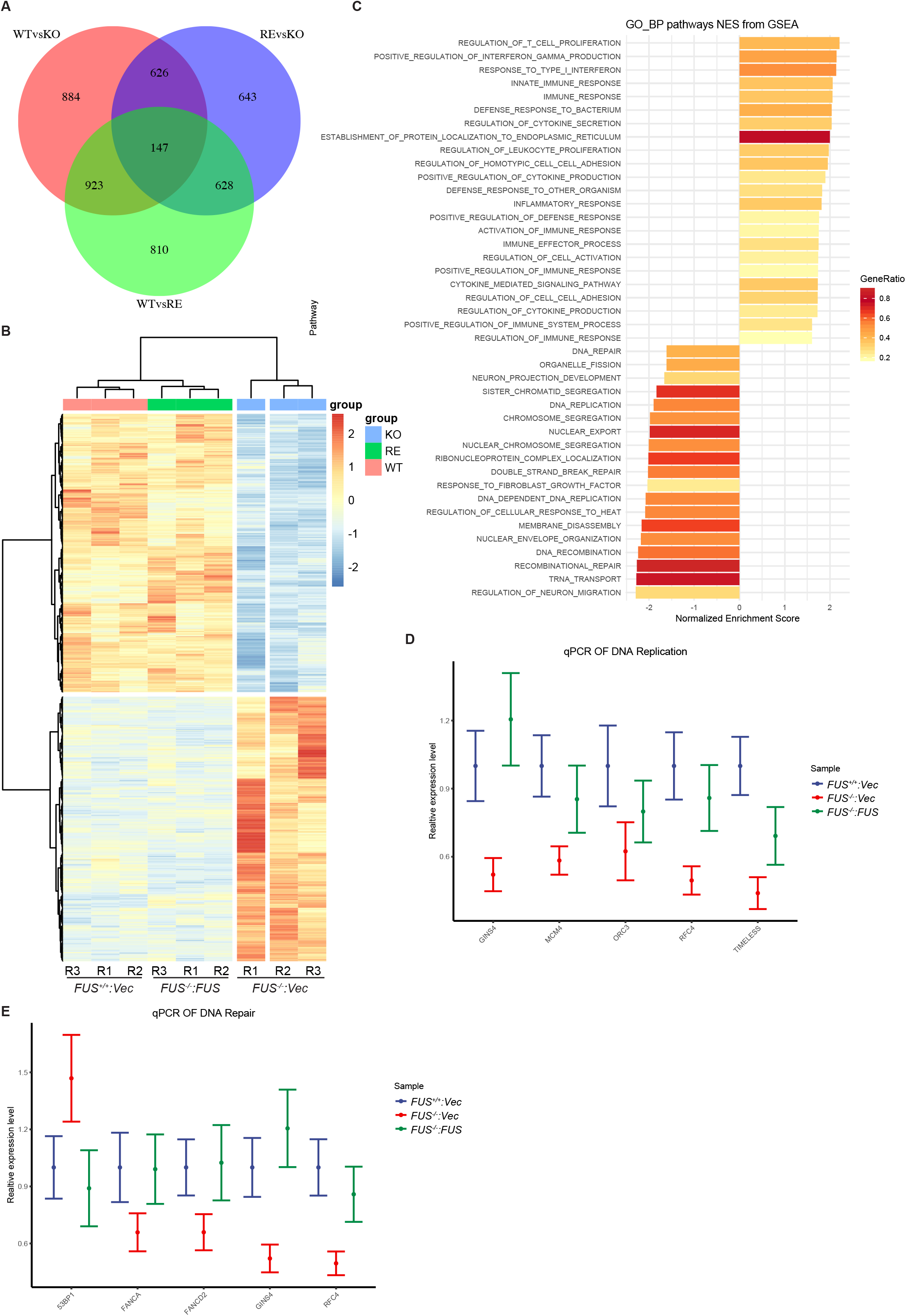
RNA-Seq and GSEA results. A, Venn diagram of differentially expressed genes (DEGs) from comparisons among *FUS^-/-^*:*Vec*(wild type, WT), *FUS^-/-^*:*Vec* (knockout, KO), and *FUS^-/-^*:*FUS* (reconstituted, RE). B, Heatmap of genes whose altered expression in *FUS^-/-^*:*Vec* cells was rescued in *FUS^-/-^*:*FUS* cells. C, GSEA results in biological processes. The GO terms shown are with adjust *p*-value of 0.01. D and E are qPCR verification results of DNA repair and replication related genes. The *β-ACTIN* and *GAPDH* genes were used as internal control normalization.

**Supplemental Figure 6.**
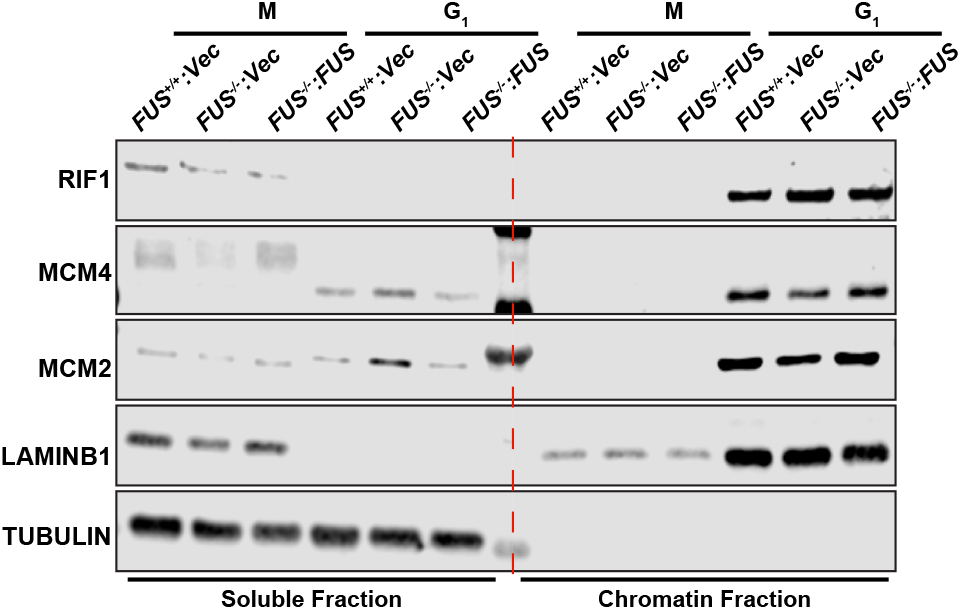
FUS regulates MCM helicase complex loading onto chromatin in G_1_ phase. Samples were prepared as shown in Fig. 5B.

**Supplemental Figure 7.**
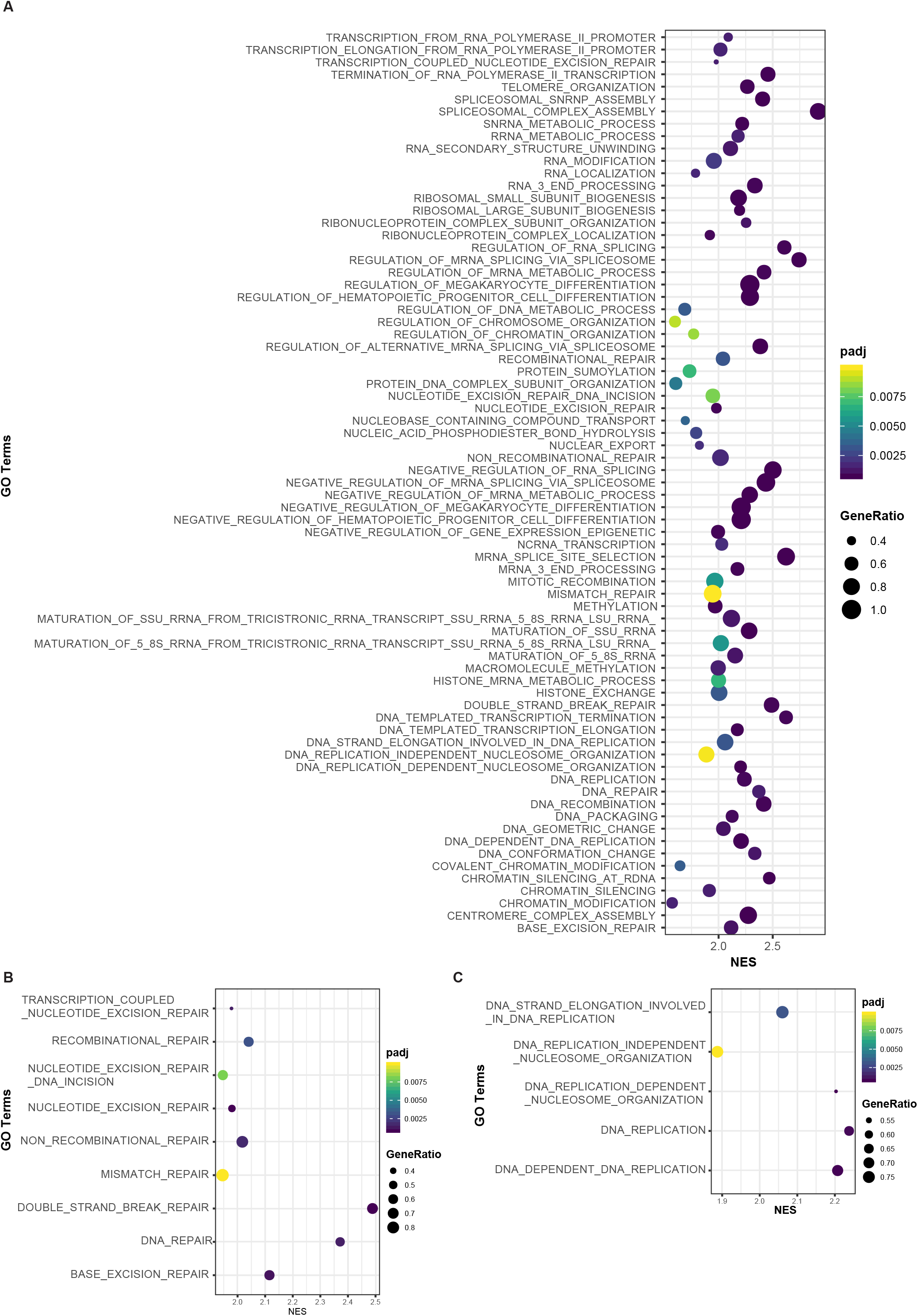
Gene set enrichment analysis of FUS interactions. A, Biological process ontology enrichment results. The enrichment was performed with R package fgsea and all pathways were shown with an adjust *p*-value lower than 0.01. DNA repair and DNA replication related enrichment pathways were shown in B and C separately.

**Supplemental Figure 8.**
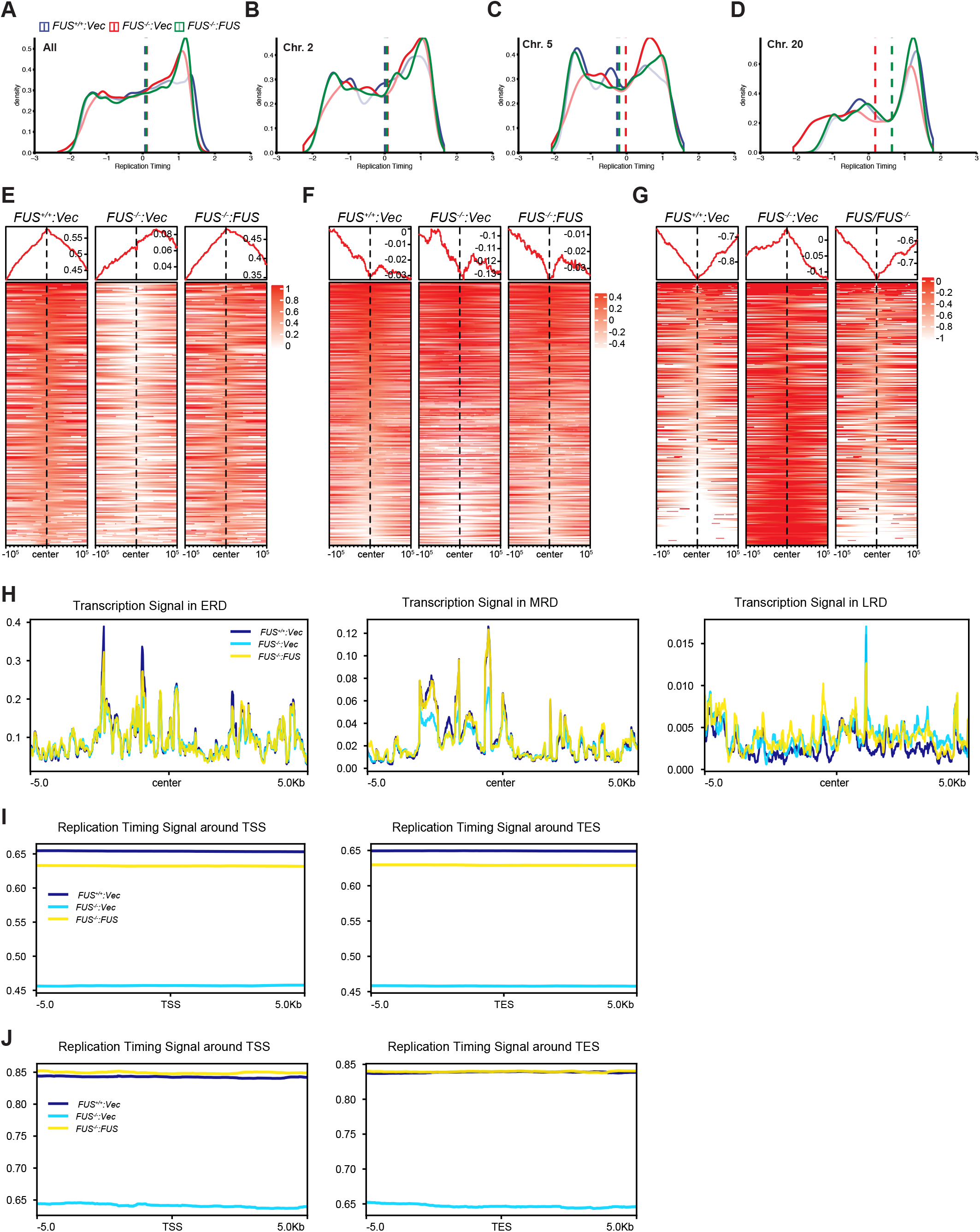
RT analysis results. A to D, the RT density distribution analysis in All chromosome(A), Chr.2(B), Chr.5(C), and Chr. 20(D) respectively in replicate 1. E to G, Heatmap results of RT signal enrichment of FAD related ERD(E), MRD(F) and LRD(G) in all individual samples, Transcription signal in the centred RDs. Transcription signal was normalized with CPM by STAR. I, Distribution of RT signal around annotated TSS and TES region across a ± 5kb window. The GENCODE v32 of GRCh38 annotation file was used. J, Distribution of RT signal around FUS regulated genes’ TSS and TES region across a ± 5kb window. Only FUS regulated genes(Sup. Tab. 4) annotation was used. The generation of RT signal was described in Fig. 9G.

**Supplemental Figure 9.**
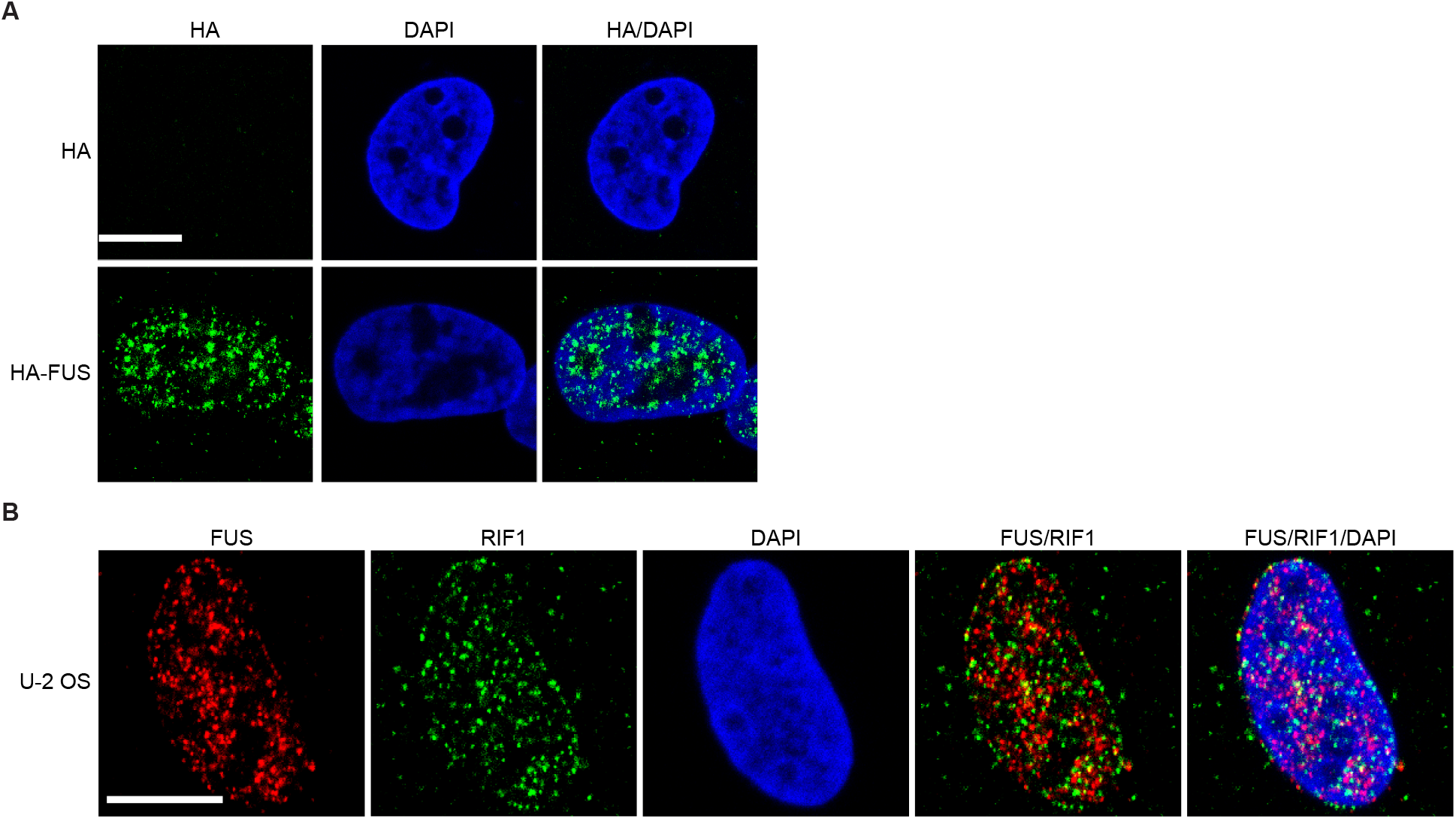
Puncta of HA-FUS in U-2 OS cells. A, Empty vector and HA-FUS plasmid were stable expressed in U-2 OS cells. Immunofluorescence imaging was preformed using α-HA HA as described in Fig. 10A. Bar size is 10 μm. B, Immunofluorescence imaging of FUS and RIF1 in cells was prepared as in A. Bar size is 10 μm.

## Supplemental Table

Table 1. list of FUS interactions identified by MS/MS.

Table 2. list of FUS interactions involved in DNA replication and repair.

Table 3. list of differentially expressed genes of FUS WT vs KO.

Table 4. list of significate expression changed genes in *FUS^-/-^*:*Vec* and rescued in *FUS^-/-^*:*FUS*.

Table 5. list of normalized counts of RNA-Seq by DESeq2

